# Disruption of Collective Behavior Correlates with Reduced Interaction Efficiency

**DOI:** 10.1101/2024.12.26.630420

**Authors:** Justine B. Nguyen, Chelsea N. Cook

## Abstract

Group-living organisms commonly engage in collective behavior to respond to an ever-changing environment. How individuals utilize local information to produce such large-scale, cohesive behaviors and the factors that impact these dynamics are of high research interest, especially since animals face increasingly challenging environmental conditions due to pollution and climate change. Western honeybees (*Apis mellifera*) are highly social insects that depend on the tight coordination of many individuals to ensure optimum colony function. We used fanning, a collective thermoregulatory behavior in honeybees that depends on both social and thermal contexts, as a case study for collective behavior. To elucidate the mechanisms behind the coordination of fanning, we used oxytetracycline, an antibiotic historically used in apiculture and known environmental pollutant that impairs bee physiology and behavior. Specifically, we hypothesized that oxytetracycline will interfere social interactions which will lead to a disruption in honeybee fanning behavior. We found that longer exposure to antibiotics decreases fanning in honeybees. Using an automated tracking software, we show that antibiotic treatment lowers the number of interactions bees have despite increasing average movement velocity, thereby impeding the social dynamics within these small bee groups. Our results contribute strong evidence that interactions between individuals may drive the collective thermoregulatory fanning response in honeybees. This work emphasizes the importance of understanding the social mechanisms that underlie critical collective animal coordination and how the effects of a common environmental pollutant on an individual can scale to affect populations.

## Introduction

Organisms across taxa utilize social and ecological information to engage in collective behavior as a ubiquitous strategy to respond to a constantly changing environment [1–4]. The regulation of schooling behavior and foraging in stickleback fish (*Gasterosteus aculeatus*), for example, largely relies on movement cues of conspecifics around them [5–8]. Harvester ants (*Pogonomyrmex barbatus*) adjust their foraging activity based on antennal cuticular hydrocarbon signals of other foraging workers that indicate environmental conditions [9,10]. Collective behavior even plays a fundamental role in regulating bacterial inter- and intra-species dynamics [11–13]. Participating in collective behavior may position social animals to deal with challenges that arise in the environment because collective behavior often allows for animals to accomplish tasks that could not be possible for individuals to perform on their own [14–16]. Understanding the factors that modulate such complex, cohesive behaviors is essential to fully comprehending how organisms interact with and respond to their environment, especially given that animals face increasing environmental challenges due to human activity and pollution [17,18].

A key aspect to the successful coordination of collective behavior is the exchange of information between group members, which can occur during social interactions [7,10,17,19–21]. Information transfer is critical in collective behaviors as it increases all participating individuals’ awareness of their environment, allows for flexibility in their decision-making process, and ultimately leads to adjustments in group-level behavior [1,22,23]. For instance, some species of ants and bees select new nest sites using only a few scouts that go on to recruit others; eventually, the entire colony coalesces to a single site, even if not all individuals have chosen the site as ideal themselves [22,24,25]. Collective foraging often depends on communication to other individuals about a resource site of interest, such as through tandem runs in ants [26,27] or the waggle dance in honeybees [28,29].

Interactions between individuals, whether direct or indirect, are often critical achieve a cohesive collective dynamic but there often exists trade-offs associated with group communication, such as sacrificing accuracy of information transfer for speed of decision-making [30–33]. Depending on the environmental context, these trade-offs could potentially result in changes in the efficiency of the interactions. In some foraging decisions, specific interaction rates can influence a participating individual’s decision to start foraging [9,34,35], but this rate can be further impacted by individual variation in movement patterns [9], response thresholds [36–38], and physiological state [33,39,40]. On the other hand, as a result of a species’ ecology, a single interaction may not be sufficient to elicit foraging; rather, a sustained series of interactions are necessary to trigger an individual to forage, as seen in harvester ants [41]. In schooling fish, the alteration of sensory perception of the presence of others via antibiotic destruction of lateral line hair cells alone disrupts social interactions [42,43]. Thus, any disruption in interaction dynamics between and across individuals, such as the disturbance of the efficiency of such interactions through physiological perturbation from environmental pollutants, could scale to affect group-level outcomes and can be used as a method to understand the factors that govern collective behavior.

Western honeybees (*Apis mellifera*) are frequently used as models of collective behavior [44–46] and are important agricultural pollinators [47,48]. Honeybees engage in various collective behaviors to respond to their environment: one of which is regulating their colony temperature within 35°C ± 1°C, which ensures proper development of their brood [49]. To keep their colonies cool, middle-aged honeybees gather at the entrance of their hive and fan their wings to circulate air into the colony, a behavior known as fanning [50]. Fanning depends on both temperature and social contexts, as bees experiencing high temperatures will rarely fan when isolated but will fan in small groups [51]. Additionally, the presence of an experienced fanner bee can influence the likelihood of a non-experienced bee to fan [52], further indicating that social interactions may be playing a role in facilitating group fanning behavior. Although we are beginning to understand the contexts and cues that are important in triggering fanning, we do not know the mechanism by which fanners work together to enact essential thermoregulation, or how this critical behavior could be potentially disrupted by environmental factors.

Here, we aimed to perturb social interactions in honeybees by feeding them an apiculturally relevant antibiotic, oxytetracycline. Historically, oxytetracycline has been the only effective method to treat honeybee colonies prophylactically and in response to foulbrood diseases [53]. However, oxytetracycline has more recently been regulated by the Food and Drug Administration (FDA) since it impacts honeybee physiology and behavior [54–56] and is a known environmental pollutant [57]. It is unknown whether oxytetracycline specifically affects interactions. We therefore hypothesized that oxytetracycline would interfere with social interactions in honeybees which would then disrupt the fanning response. We specifically predicted that oxytetracycline treatment would decrease the number of bees that fan in response to increasing temperatures in a group and increase the temperature threshold at which they begin to fan. We also predicted that the decrease in fanning performance in treated bees would be a result of decreased number of interactions. By exploring this hypothesis, we provide insight into how honeybees socially organize and thermally manage their environment and how exposure to an environmental pollutant could potentially disrupt these dynamics.

## Methods

All experiments took place between May and October 2023. Honeybees (*Apis mellifera l.*) used in experiments described below originated from managed colonies located on the roof of Wehr Life Sciences at Marquette University in Milwaukee, Wisconsin. Colonies were maintained using best practices ensuring they kept healthy weights and population sizes. All colonies were monitored for disease and treated for Varroa mites using ApiGuard (thymol) following label instructions and oxalic acid vaporization twice during the experimental period: once in August and again in September. We collected bees from 8 of these managed colonies.

### Fanner collection

To identify fanners, we collected bees on the porch of their colonies who were standing with their heads angled towards the entrance of their colony and actively fanning still for at least 10 seconds [51]. To distinguish fanners from bees who were Nasanov fanning as a homing signal, we chose bees who were fanning with their abdomens angled downwards without their Nasanov’s gland exposed [51]. For our study, we did not explicitly control for age, though generally, fanners tend to belong to the middle-aged task group with a mean age of 14.7-19 days old [50,58,59]. Although fanning occurs throughout the colony, we collected fanners from their colony entrance as they are easily accessible and have been previously demonstrated to be relatively uniform in age [50,51]. Fanners were collected using flexible, soft forceps into empty pipette tip boxes with 96 well plates taped inside to mimic natural comb structure. To minimize light and thermal stress during collection, all boxes were kept in the shade once collected. Fanners were placed together in plastic boxes (4.5in x 3.25in x 2.5in) in groups of 30, and each box consisted of fanners collected from the same colony. Three boxes of fanners (one box per treatment group as later described) were collected per colony. Depending on fanner availability, up to two more sets of triplet boxes were collected from other colonies on the same day.

### Experimental treatment groups

After collection, all boxes were brought back into the lab and were randomly assigned to one of 3 experimental treatment groups: sugar syrup solution only (referred to as “lab control”), sugar syrup solution for 4 days then 24 hours of sugar solution with antibiotic (referred to as “1-day treatment”), or sugar solution with antibiotic for 5 days (referred to as “5-day treatment”). We assigned treatments where all three treatment groups were from the same colony and repeated these assignments throughout the experiment across 8 colonies. Bees were provided an *ad libitum* source of their treatment solution. Sugar syrup solution was 40% concentration, while antibiotic treatment was made as a 40% sugar syrup solution with 1mg/mL of the antibiotic oxytetracycline hydrochloride (Oxytet® Soluble), procured through a local veterinarian. FDA recommendations for the administration of oxytetracycline (200mg) as of 2020 focuses on the treatment of the whole hive [60] and is therefore difficult to scale to what individual honeybees may experience. We aimed to utilize a physiologically disruptive dose in our study. Previous studies identified 1 mg/mL to be toxic to bee larva [61,62] and 0.45mg/mL was used in studies looking at microbiome disruption of antibiotics (0.45mg/mL) [54]. Oxytet® Soluble has 37% oxytetracycline, so using 1 mg/mL of product gave a treatment of 0.37mg/mL of active ingredient and is similar to previous studies [54]. Boxes with bees fed their respective treatments were kept in a dark incubator at 35°C with 40% relative humidity for a total treatment period of 5 days in accordance with other honeybee antibiotic studies [54]. Dead bees were removed and counted daily for survival data.

### Fanning assay

To test fanning capabilities after the treatment period, all surviving fanners were subjected to a well-established fanning assay [51]. Briefly, we transferred groups of 5 experimental bees into wooden framed wire mesh cages (340.47 cm^3^) from the same colony and same treatment. Cages were then placed inside large glass jars on top of hotplates (OHause® Guardian™5000 Model number e-G51HP07C) with a temperature probe (Campbell Scientific, 109 Temperature Probe) wired through a GRANITE™ Volt 116 16-/32-Channel 5V analog input module connected to a computer as our data acquisition system. SURVEYOR software (Campbell Scientific, v. 1.01) was used to monitor and collect data on the temperature inside the jars in real time. Bees were allowed to acclimate at room temperature (mean temperature ± standard deviation, 23.63 ± 0.93) for 25 minutes before the temperature inside of jars was continuously increased by experimenters by 1°C per minute via the hot plate. Bees were then observed for fanning behavior using the same behavioral metrics as collecting fanners (i.e., standing still for at least 10 seconds, abdomen curved downward, and actively flapping wings [51]). The time, temperature, and number of fanners seen fanning throughout the assay were recorded [51]. Fanning frequently occurs in ‘bouts’, where small groups of bees will begin fanning within a minute of each other, so we recorded the initial number of bees fanning within a minute of each other as the ‘first bout” (i.e., the first bout of fanning in the trial), and any additional fanners that joined later were counted within ‘max bout’ (i.e., the bout where the maximum number of bees were simultaneously fanning during the trial). First bout and max bout can be the same bout which occurred in 38 of the trials. Bees were observed until mortality. To reduce bias in recording observation data, experimenters were blinded to the identity of each fanning group before cages were placed in jars. After trials concluded, we calculated actual trial ramp rates and removed any trials that were not within 0.1°C of 1°C/min from our analysis. We also collected a group of fanners immediately before fanning trials (referred to as ‘hive control’) to serve as a behavioral comparison to hive bees.

### Video Recording

See Supplemental Figure 2 for a photo reference and description of our video recording set up. To measure social interactions within fanner groups across treatments, we built a video recording station (Supplemental Figure 2A) and an experimental apparatus (Supplemental Figure 2B) of similar volume to fanning cages (354.98 cm^3^). After treatment, fanners were marked on their thorax with different colors using a non-toxic water-based paint marker before being transferred to the video recording box. Like in regular fanning assays, fanners were placed in the video recording box (Supplemental Figure 2B) in groups of 5 of the same treatment and from the same colony. The wooden box was then placed into a 13 x 8.5 x 3.25-inch glass baking dish (Pyrex) with a glass lid (Supplemental Figure 2C) on top of a hotplate (Supplemental Figure 2D) to mimic the well-established fanning assay protocol. Bees were allowed to acclimate inside of the recording set up for 25 minutes, and experimenters began video recording using a Panasonic HC-V800 camera placed above the station through a hole 5 minutes (Supplemental Figure 2E) before the trial started. Videos were filmed at a 30 frames per second resolution. A computer monitor (Supplemental Figure 2F) was connected to the camera to allow for experimenters to observe the bees. Once the trial started, hot plates were ramped at 1°C per minute as in a usual fanning trial and experimenters watched the computer monitor for behavior. We recorded the number of fanners, the colors of the bees who fanned, time, and temperature as described previously, including first fanning bouts and max fanning bouts. Once the fanning trial concluded, experimenters immediately stopped video recording and calculated ramp rates. Any trials that were not within 0.1°C were not used in further analysis. All videos were backed up on an external hard drive and transferred to a computer for further tracking analysis.

### ABCTracker

To extract data from our videos, we used the tracking computer program ABCTracker [63,64]. To prep our videos for the ABCTracker pipeline, we identified when the first instance of fanning occurred using observation data within the video and created 5-minute trims: 4 minutes before the first instance of fanning and 1 minute after. This allowed us to account for normal variation in acclimation and initial fanning temperature across groups. We repeated this process until we had fanning trims from all videos. Processed data videos were backed up on Marquette University’s cloud-based Microsoft OneDrive service for long-term storage.

We then uploaded fanning video trims in .mp4 format into ABCTracker, marked all five bees as trackable objects, and allowed the program to run its automatic tracking process. Then, an experimenter manually corrected any errors the program made in tracking individuals throughout the trim. Once video tracking was complete, .csv files with raw tracking data were downloaded for analysis. These files, which included x-y-coordinates and velocity (pixels/second) of each bee per frame, were then fed into an R Studio pipeline that cleaned, extracted, calculated the necessary data of interest to our study. This pipeline detects head-to-head, head-to-body, and body-to-body interactions by calculating when and where we first identify an interaction as an overlap between the x-y coordinates of the heads, bodies, or heads and bodies of two bees that lasted for at least a second, or the equivalent of 30 frames of video [65]. Multiple interactions could occur at the same time, so we established an interaction hierarchy where head-to-head was the primary interaction, then head-to-body, then body-to-body. For example, if head-to-head and body-to-body happens within the same interaction, we classify it as head-to-head. When we plotted interaction type as a proportion of all interactions the bees had within a video, head-to-head interactions accounted for nearly 75% of all interactions across all treatment groups (Supplemental Figure 1). We therefore chose to focus on head-to-head interactions and filtered all other types of interactions out of our tracking data.

### Statistical Analysis

All data analyses were performed using R (v4.3.2) and R Studio (v4.2.0). We used the following R packages for our analysis: “car”, “data.table”, “dplyr”, “emmeans”, “fitdistrplus”, “FSA”, “ggplot2”, “ggpubr”, “lme4”, “patchwork”, “pastecs”, “performance”, “reshape”, “tibble”, and “tidyverse”.

To test for statistical differences in percent survival after treatment duration, we first calculated the proportion of bees that survived their treatment. We then used a generalized linear mixed model to create a logistic regression. After verifying model assumptions and running appropriate diagnostics (i.e. normality of residuals, homoscedasticity, independence of errors, etc.), we then tested the model via an ANOVA type II test which was followed by a Tukey post-hoc for pairwise comparisons. Our predictor variable was treatment while our response variable was percent survival. We calculated survival likelihood by extracting and exponentiating coefficients from our model and calculating probabilities from the resulting odds ratios. We then generated our survival curves using ggplot2.

To test for statistical differences in our behavioral fanning data, we created several generalized linear mixed models (GLMM), verifying model assumptions before continuing the analysis. First, we calculated the proportion of max fanning bout fanners and first fanning bout fanners by dividing the number of fanners by the total number of bees tested in the fanning trial. We then created logistic regressions via GLMM using these proportions. We tested the model via an ANOVA type II test to determine if the main treatment effects were significant and then used a Tukey post-hoc test for pairwise comparisons when appropriate. Our predictor variable was treatment, and our response variable was proportion of fanners out of the total number of bees in the group. We calculated likelihood of fanning using the same conversion method as calculating survival likelihood. For all temperature threshold data, we created a linear regression model (LM) followed by an ANOVA type II test. Our predictor variable was treatment, and our response variable was the temperature at which bees first began to fan and the temperature at which the maximum number of bees fanned. To test for statistical differences in temperatures at first observed death, we created a linear regression model, followed by an ANOVA type II test and a Tukey post-hoc test. Our predictor variable was treatment, and our response variable was the temperature at which the first death was observed. Because we collected fanners from 8 different colonies, colony was originally incorporated as a random effect in all our models. However, colony origin accounted for minimal variation across all our models (treatment survival model, variance = 0; first fanning bout temperature threshold model, variance = 0.4139 ± 06434; max fanning bout temperature threshold model, variance = 0.9829 ± 0.9914; proportion of fanners in the first bout model; variance = 0, proportion of fanners in the max bout model; variance = 7.438 x 10^-14^ ± 2.727 x 10^-7^); as a result, we removed colony as a random effect from all our models and proceeded with using non-mixed models.

To test for differences in movement velocity and number of interactions across treatment groups with our video tracking data, we quantified the number of head-to-head interactions and compared them across treatment groups using a Kruskal-Wallis test and Dunn’s test for post-hoc analysis, as data violated normality and homogeneity of variance assumption tests (Shapiro-Wilk and Bartlett’s test, respectively). To calculate average group velocity, we calculated the total distance bees moved, divided it per video frame, and then took the average velocity per frame. We then created linear regression models with our predictor variable as treatment, and our response variable as average group velocity. Because time and temperature are colinear in our analysis, as our ramp rates per video were consistent at 1°C/min, by using velocity as our response variable, our model accounted for time. See Supplemental Table 1 for information of our model estimates. To analyze any patterns in our interaction data, we also created a GLM with our predictor variable as treatment and our response variable as number of head-to-head interactions. We tested our models for statistical differences via an ANOVA type II test and then used a Tukey post-hoc test for pairwise comparisons.

Based on the previous results on movement velocity and number of interactions, we calculated an interaction efficiency index to better understand the role of these metrics in the social dynamics of our groups of bees. To calculate an interaction efficiency index, we divided the number of head-to-head interactions by average group velocity at each time point. Because efficiency is classically defined as output over input, we chose to calculate our efficiency index with average group velocity as our ‘input’ and number of interactions as our ‘output’. Our rationale was that honeybees would have to move around the video arena (input) to access other bees and participate in interactions (output). To test for differences in social efficiency, we used a linear regression model with efficiency as our response variable and treatment as our predictor variable, testing model assumptions before continuing with the analysis. An ANOVA type II test was then used to test the model, followed by a Tukey post-hoc test for pairwise comparisons.

## Results

### 5-day exposure to antibiotics disrupts fanning behavior

We found that 5-day treatment bees were less likely to fan than 1-day or control treatment bees during both the first bout and max bout of fanning seen in trials (Table 1). This led to a smaller proportion of 5-day treatment bee groups fanning during the max fanning bout (Figure 1C, X^2^ = 15.695, p = 0.001). Although we detected a statistically significant difference between groups within our first fanning bout data (Figure 1A, X^2^= 7.85, p = 0.0492), our post-hoc tests could not detect which groups were statistically different from each other. Additionally, we found that 5-day antibiotic exposure decreases the ability of honeybees to withstand high temperatures, as 5-day treatment bees began to die at lower temperatures on average (5-day treatment mean ± standard deviation death temperature, 46.24°C ± 7.53) during fanning assays compared to all other groups (Figure 2; X^2^= 17.04, p = 0.001, Tukey’s post-hoc test, p < 0.01). These results replicated our previous findings (See Supplementary Figures 3 & 4). 1-day treatment bees performed fanning just as well as hive control and lab control bees (Figure 1; Tukey’s post-hoc test, p > 0.05), and there was no difference in the temperature at which they died (Figure 2; Tukey’s post-hoc test, p = 0.6).

**Table 1.** 5-day exposure to antibiotics decreases likelihood of fanning. Lab control = bees exposed to a 40% sucrose solution for 5 days, 1-day treatment = bees exposed to oxytetracycline 24 hours prior to fanning assay, and 5-day treatment = bees exposed to oxytetracycline for 5 days, hive control = regular fanners collected directly from colonies the day of testing. We defined the initial number of bees fanning within a minute of each other as ‘first bout’, and any additional fanners that joined later and were part of the maximum number of fanners seen to be fanning in the trial were counted within the ‘max bout’ of fanning. We calculated fanning likelihood by extracting and exponentiating fixed effect coefficients from our logistic regression model and calculating probabilities from the resulting odds ratios.

| Treatment Group | Likelihood of Fanning in First Bout | Likelihood of Fanning in Max Bout |
| --- | --- | --- |
| Lab control | 0.303 | 0.636 |
| 1-day | 0.242 | 0.564 |
| 5-day | 0.206 | 0.419 |
| Hive control | 0.338 | 0.538 |
| **Test statistic and p-value (ANOVA type II test)** | $X^2 = 7.726$, $p = 0.05$ | $X^2 = 15.363$, $p = 0.002$ |

**Figure 1.**
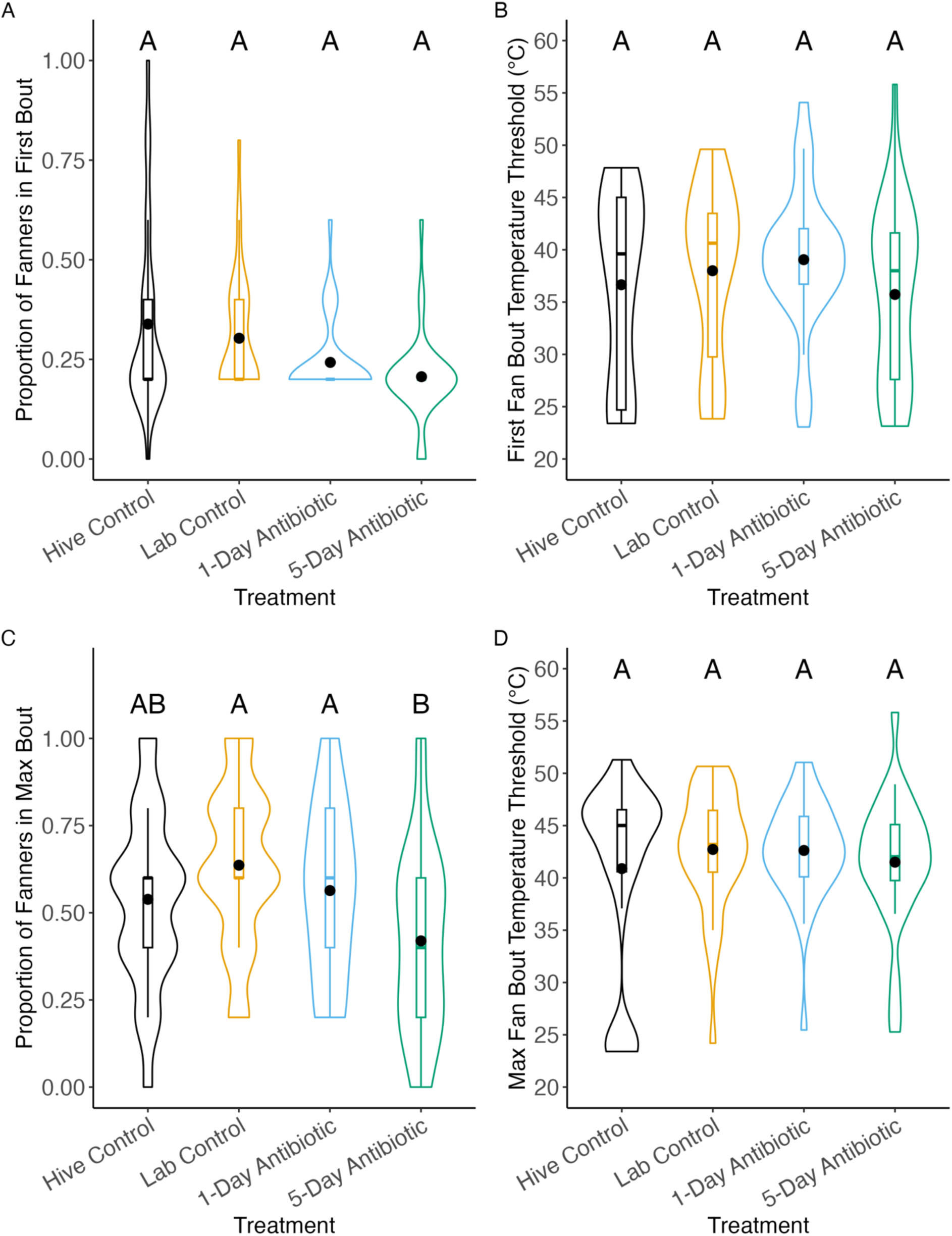
5-day exposure to oxytetracycline reduces fanning performance but not thermal threshold in honeybees. Fanner honeybees subjected to oxytetracycline treatment experience a deficit in their fanning behavior in comparison to bees not treated with antibiotics. These effects do not extend to the fanners’ temperature thresholds. Hive control = fanner bees collected from the entrance of colonies the day of assay, lab control = bees exposed to a 40% sucrose solution for 5 days, 1-day = bees exposed to oxytetracycline 24 hours prior to fanning assay, and 5-day = bees exposed to oxytetracycline for 5 days. n_hive control_=26, n_lab control_ = 33, n_1-day_ = 33, n_5-day_ = 31, for n = cage of 5 bees. Boxplot middle lines indicate median, while middle black dots indicate mean. Letters on boxplots indicate statistical difference. A) While there were no significant statistical differences detected between groups, there was a trend towards treatment bees having a smaller proportion of fanners in the first bout. Logistic regression, X^2^ = 7.85, p=0.0492 B) There were no differences in the temperature threshold of the first bout. Linear regression, X^2^ = 3.0291, p=0.39. C) Honeybees that were treated with oxytetracycline for 5-days are significantly less likely to fan than the lab controls and 1-day treatment bees. Logistic regression, X^2^ = 15.363, p = 0.002, Tukey post-hoc test, p<0.05 D) There were no differences in temperature threshold during the max bout of fanning. Linear regression, X^2^ = 1.7898, p=0.62.

**Figure 2.**
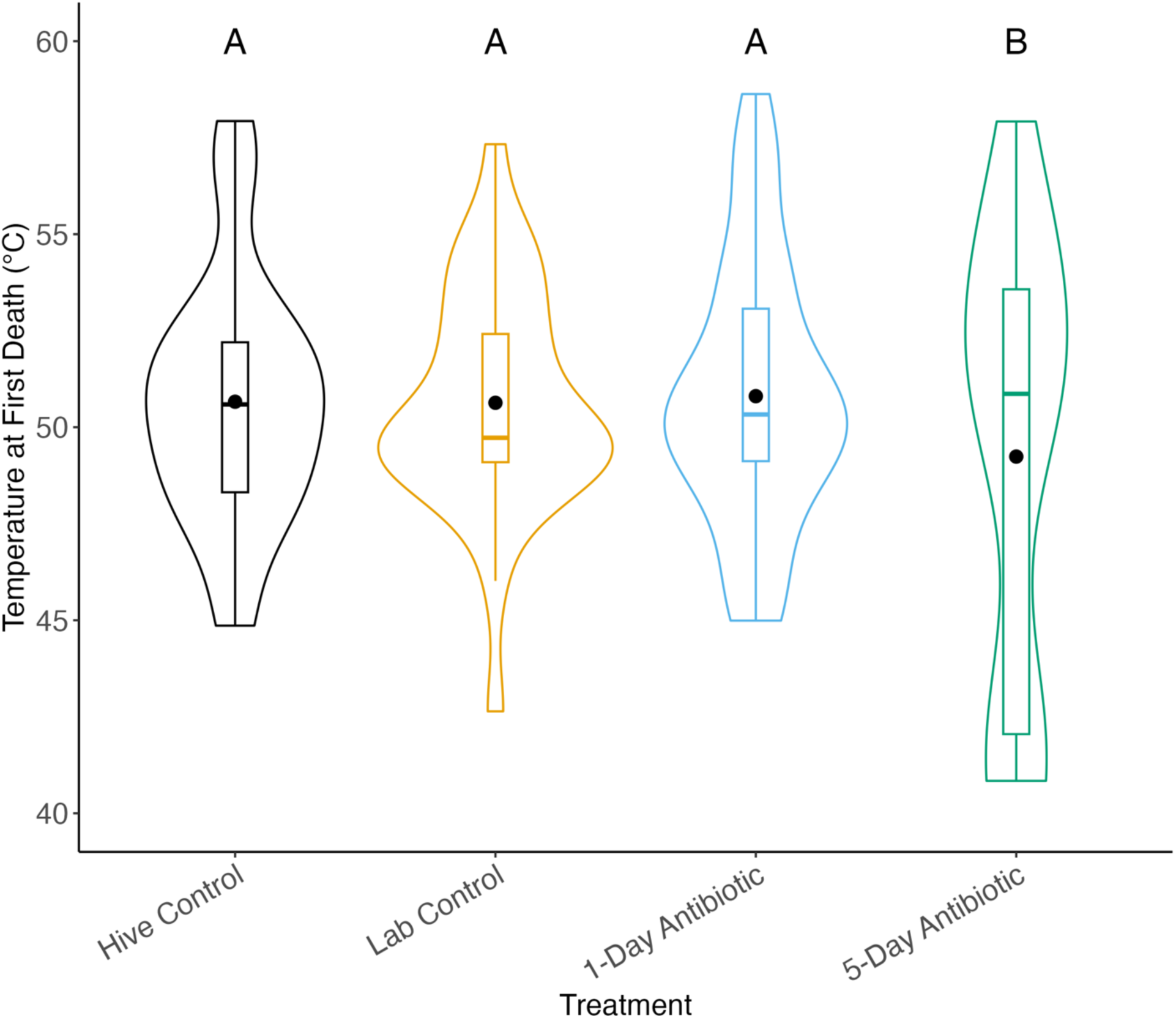
5-day antibiotic treatment lowers the temperature at which the first death occurs during fanning assay. Boxplot middle lines indicate median while black middle dots indicate mean. n_hive control_=26, n_lab control_ = 33, n_1-day_ = 33, n_5-day_ = 31, for n = number of cages of 5 fanner bees. Letters on boxplots indicate statistical difference. Linear regression, F= 6.0976, p = 0.001, Tukey post-hoc test, p < 0.05.

### 5-day antibiotic treatment decreases the number of social interactions but increases average group velocity

We found that groups of 5-day treatment bees consistently moved at high velocity (mean velocity ± SE, 364 ± 9.79 pixels/second) throughout the duration of video fanning trials (Figure 3A & C; F = 84.809, p < 0.001; Supplemental Table 1). This result contrasted with all other video recorded groups, which showed velocity significantly slowing down in the seconds leading up to the first instance of fanning (Figure 3C; Supplemental Table 1). We also began to see antibiotic effects in 1-day treatment bees (Figure 3C & D; X^2^ = 23.973, p < 0.001, Dunn’s post hoc test, p <0.001; X^2^ = 124.13, p < 0.001; Dunn’s post hoc test, p < 0.001), as they also moved at on average higher velocities (mean velocity ± SE, 245 ± 11 pixels/second) compared to lab control bees (mean velocity ± SE, 161 ± 6.3 pixels/second) and hive control bees (mean velocity ± SE, 238 ± 9.64 pixels/second) and did not have as many interactions (mean number of interactions ± SE, 108 ± 24) as lab control bees (mean number of interactions ± SE, 167 ± 36.1). However, although 5-day treatment bees on average moved at high velocity, we found that this did not translate to more interactions (mean number of interactions ± SE, 110 ± 19.5) compared to all other groups (Figure 3B & D). Counterintuitively, we found that hive control bees also moved at high velocity (mean velocity ± SE, 238 ± 9.64 pixels/second) but had lower number of interactions (mean number of interactions ± SE, 135 ± 21.5) compared to lab control (Figure 3).

**Figure 3.**
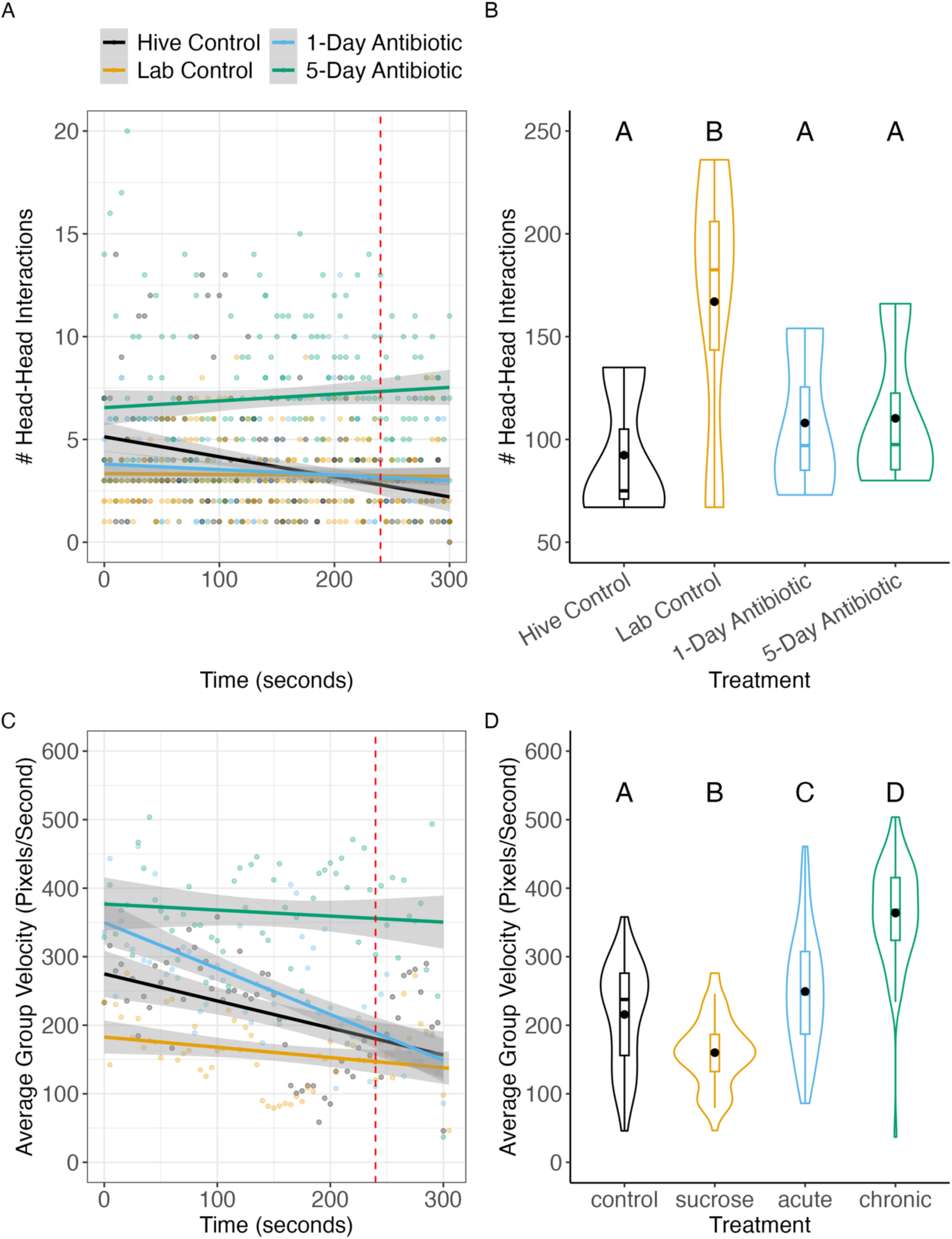
Antibiotics increase average group velocity and decrease number of interactions the group has over time. A) Number of head-to-head interactions treatment groups had over the duration of the assay. Shaded regions indicate 95% confidence intervals Each dot on line graphs represents average number of interactions the group had at a specific time for a given video replicate of a treatment group. Linear regression, F = 88.123, p < 0.001. B) Number of head-to-head teractions across experimental groups. Kruskal-Wallis Test, X^2^ = 23.973, p < 0.001; Dunn’s post hoc test, p <0.001 C) Average group velocity (pixels/second) was calculated using x-y-coordinate data from ABCTracker. Each dot on line graphs represents average group velocity for a given video replicate of a treatment group every 10 seconds. Linear model, F = 84.809, p < 0.001. Trend lines were made using a linear regression method. D) Average group velocity in pixels/second across experimental groups. Kruskal-Wallis Test, X^2^ = 124.13, p < 0.001; Dunn’s post hoc test, p < 0.001. n_hive control_ = 3, n_lab control_ = 4, n_1-day_ = 3, n_5-day_ = 4, for n = number of videos recording a group of 5 bees. Red dotted lines at 240 seconds on line graphs indicate time when first instance of fanning occurs across all videos. Letters on boxplots indicate statistical difference, black dots on boxplots indicate mean, and middle lines on boxplots indicate median.

### 5-day exposure to antibiotics decreases interaction efficiency

We found that 5-day treatment bees are less efficient (mean efficiency ± SE, 0.0264 ± 0.001) in their social dynamics and do not change their efficiency over time as temperature increases (Figure 4). This is likely because they appear to invest in movement over having interactions (Figure 3). On the other hand, lab control, 1-day treatment, and hive control bees increased their efficiency leading up to the first instance of fanning in videos (Figure 4A). This is in line with what we saw in our velocity and interaction data (Figure 3). Hive control bees were less efficient (mean efficiency ± SE, 0.0395 ± 0.00261) in their social interactions compared to lab control bees (mean efficiency ± SE, 0.0552 ± 0.00227), despite being able to perform fanning at a similar level (Figure 4B). 1-day treatment bees also were less efficient (mean efficiency ± SE, 0.0389 ± 0.00193) than lab control bees (Figure 4B).

**Figure 4.**
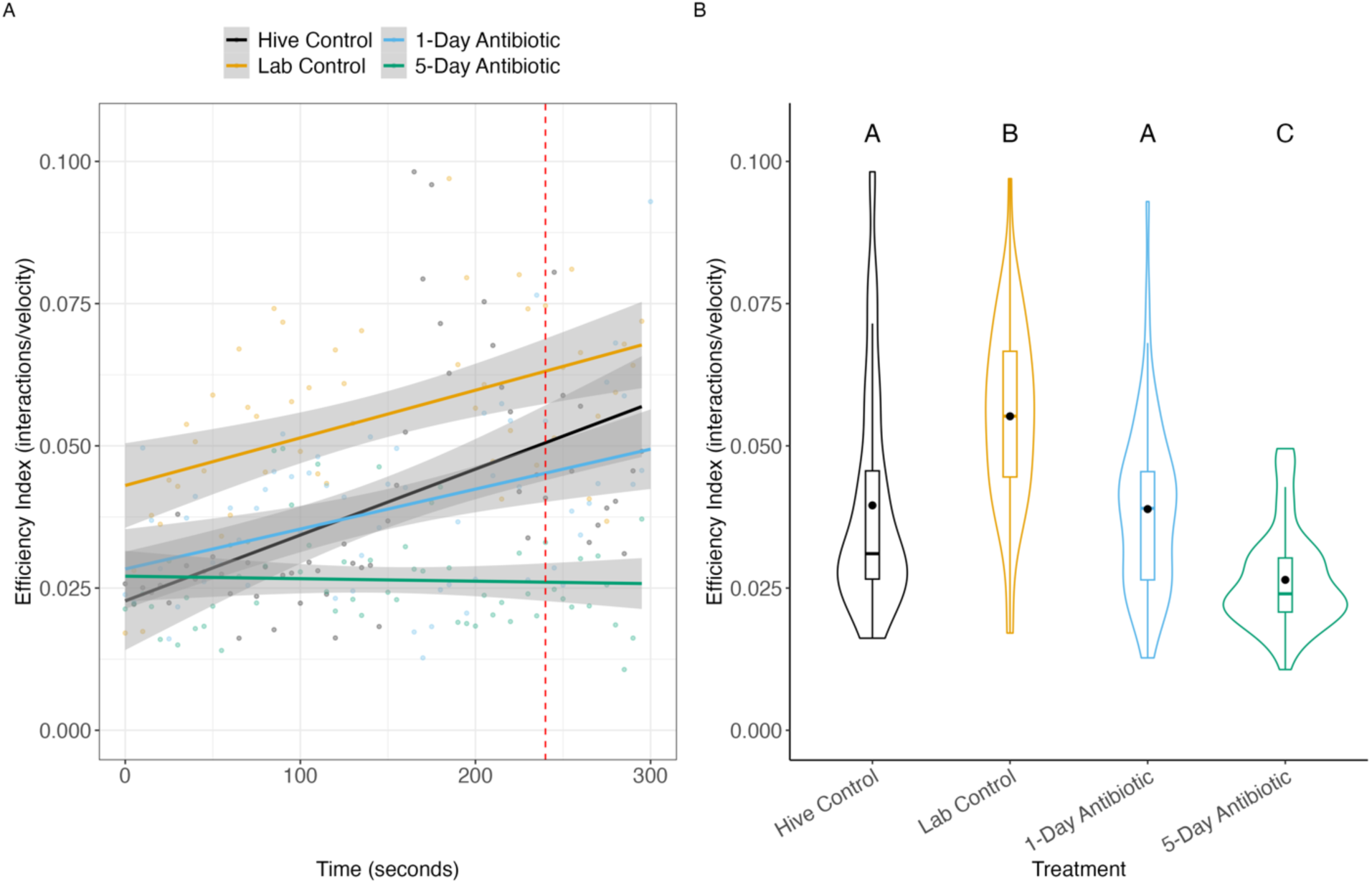
Antibiotics decrease interaction efficiency. n_hive control_ = 3, n_lab control_ = 4, n_1-day_ = 3, n_1-day_ = 4, for n = number of videos recording a group of 5 bees. We calculated an efficiency index as total number of interactions / average group velocity, which we interpreted as the ability for a group of bees to effectively communicate with each other (see Methods for how we decided on this calculation. A) Efficiency across treatment groups over time during fanning trial. Shaded regions indicate 95% confidence interval, and trend lines were created using a linear regression method. Each dot represents efficiency index of a given group replicate every 10 seconds. Red dotted line indicates time when first instance of fanning occurs. Linear regression, F = 32.35, p < 0.001. B) Average efficiency index across treatment groups. Kruskal-Wallis test, X^2^ = 83.482, p < 0.001; Dunn’s post hoc test, p < 0.001. Black dot indicates mean, while middle line indicates median. Letters indicate statistical differences.

**Table 2.**
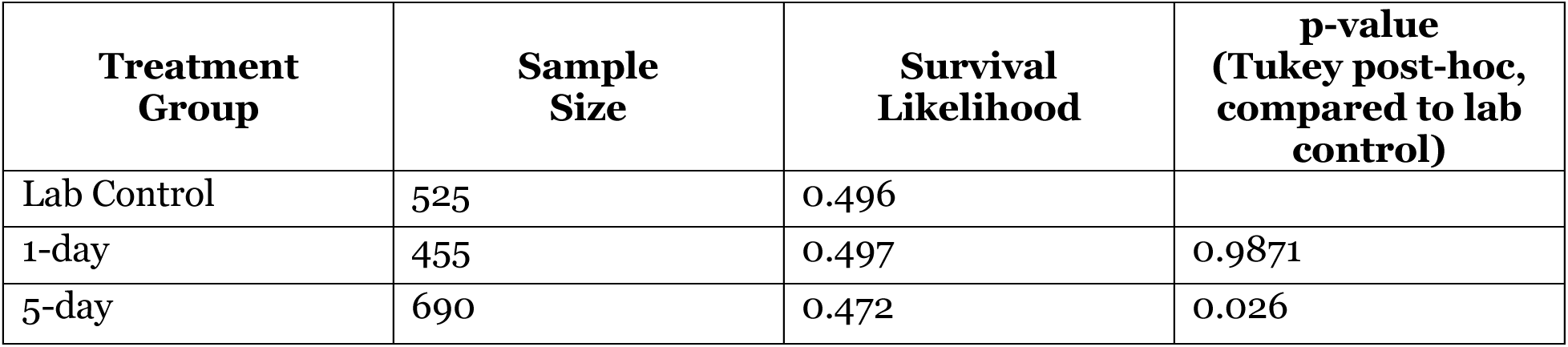
Long term exposure to antibiotics decreases treatment survival. Sample size indicates individual bees collected and subjected to treatment. Lab control = bees exposed to a 40% sucrose solution for 5 days, 1-day = bees exposed to oxytetracycline 24 hours prior to fanning assay, and 5-day = bees exposed to oxytetracycline for 5 days. p-values obtained using Tukey post-hoc analysis, comparing sucrose to all other treatment groups. Logistic regression, X^2^ = 11.988, p = 0.002, Tukey post-hoc test, p < 0.05. Survival likelihood was calculated by extracting log probabilities from our logistic regression model and calculating probabilities using the appropriate conversion.

| Treatment Group | Sample Size | Survival Likelihood | p-value (Tukey post-hoc, compared to lab control) |
| --- | --- | --- | --- |
| Lab Control | 525 | 0.496 |  |
| 1-day | 455 | 0.497 | 0.9871 |
| 5-day | 690 | 0.472 | 0.026 |

**Figure 5.**
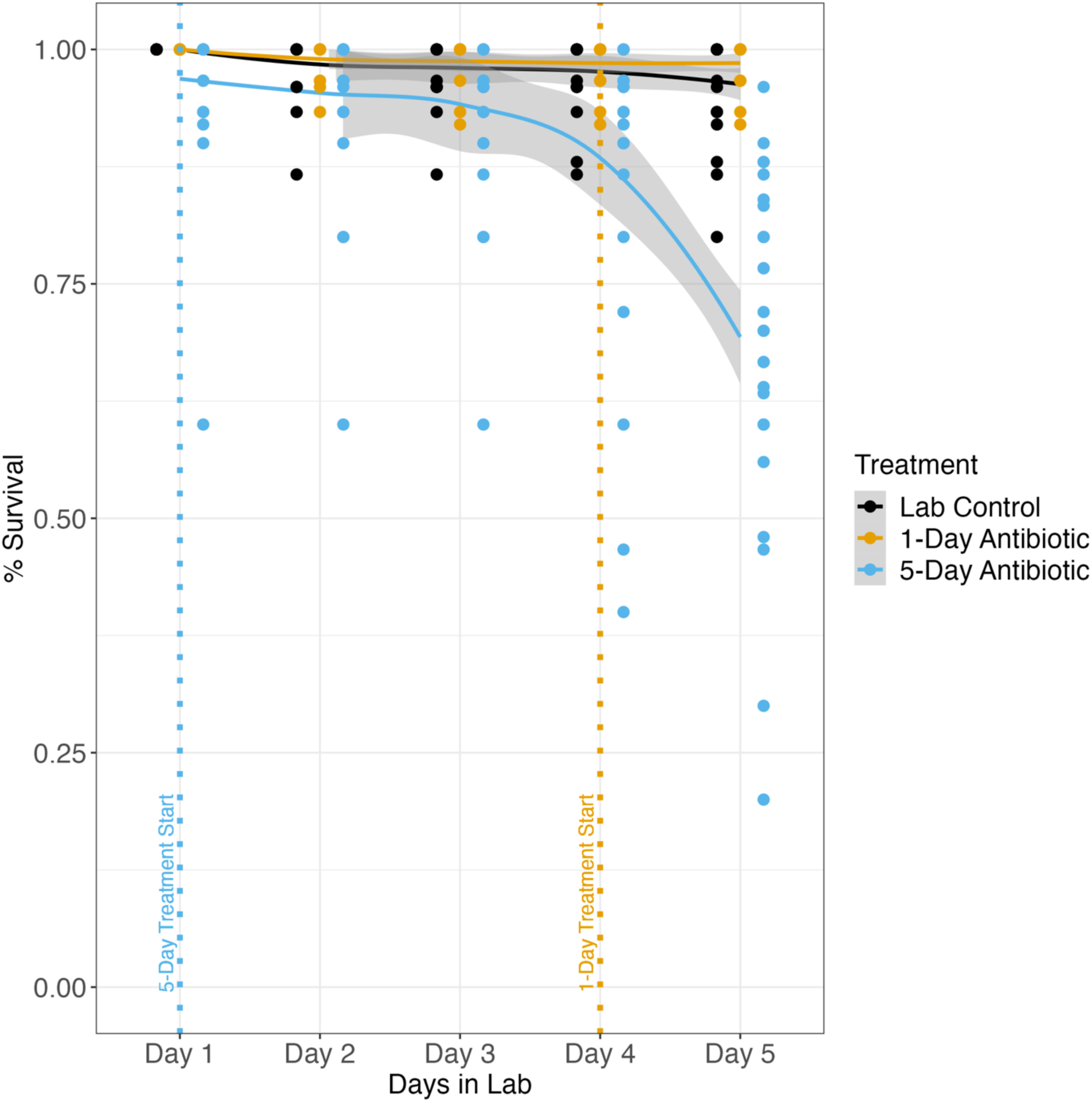
5-day antibiotic treatment decreases honeybee survival during treatment duration. Survival curves for the duration of our experimental treatment period. All fanner bees were housed in groups of 30 consisting of same colony members. Fanners that were treated for 5 days with oxytetracycline died more over the course of treatment period. Logistic regression, X^2^ = 9.5902, p = 0.008249, Tukey post-hoc test, p < 0.05. n_lab control_= 20, n_1-day_= 17, n_5-day_= 26, for n = boxes of 30 bees. Shaded regions indicate 95% confidence intervals.

### 5-day antibiotic treatment decreases survival in fanner honeybees held in lab

5-day antibiotic treatment decreased survival during the 5-day treatment period (Figure 5). More bees died during the duration of the treatment in the 5-day group compared to all other groups (Table 2; Figure 5; X^2^ = 11.988, p = 0.002, Tukey’s post-hoc test, p < 0.05). Interestingly, the number of deaths per day sharply increased starting on day 4 for the 5-day treatment bees (Figure 5). These findings replicated what we saw previously (See Supplementary Figures 2 & 3). The 1-day treatment and general lab rearing conditions had no effect on survival throughout treatment duration (Table 2; Figure 5).

## Discussion

In this study, we identified that social interactions play a role in the performance of collective fanning behavior in honeybees. By using antibiotics as a disruptor of individual physiology, we found that a 5-day exposure to antibiotics reduced group fanning response to high temperatures (Figure 1; Table 1). By tracking groups of bees as they experience increasing temperatures, we discovered that the disruption in the fanning response is associated with lower numbers of head-to-head interactions leading up to the first fanning bout (Figure 3). Interestingly, the number of interactions dropped despite higher average group velocity (Figure 3), demonstrating that there may be an efficiency trade-off in social dynamics within these small groups (Figure 4). These findings support the hypothesis that animal groups must balance the resources required (e.g., energy) for individuals to effectively work together and produce a collective behavior [30–33]. Ultimately, this balance may depend on the decision-making process of all individuals within a group [66], and overall group efficiency in balancing these trade-offs can determine the success of groups in responding to their environment.

The honeybee fanning response is characterized by both the probability of performing the behavior and the temperature threshold at which the behavior is performed [51]. In our experiment, fanning occurred at similar temperatures regardless of duration of antibiotic treatment (Figure 1). Antibiotic treatment and duration likely had little impact on thermal sensory processing for the honeybees, meaning they can still accurately sense temperature, yet they appear to be less effective in coordinating a fanning response to high temperatures. These results further support our hypothesis that social interactions are important in eliciting collective fanning behavior in honeybees.

Survival during the treatment period and during the fanning assay was impacted by duration of antibiotic treatment. An antibiotic exposure period of 1-day did not impact treatment survival, while a 5-day antibiotic exposure significantly increased mortality during treatment duration (Figure 5; Table 2). However, 1-day treatment did begin to mildly affect bee social dynamics; preceding the first instance of fanning, these bees had fewer interactions, higher average velocity, and therefore lower efficiency relative to lab control bees (Figure 3 & 4). While 1-day treatment may not be enough to entirely disrupt fanning performance, tolerance to high temperatures, or survival, it is notable that oxytetracycline has been demonstrated to persist in colonies for several months after antibiotic treatment [67]. Furthermore, the temperature of first recorded death was lower in 5-day treatment bees compared to all other groups (Figure 2). The heat shock response and sociality are linked: social interactions have been implicated in the induction of the heat shock response, suggesting that social environment may potentially play a role in either mitigating or exacerbating stress [68–70]. Hive control bees also had fewer interactions and were less efficient in their social interactions than expected (Figure 3 & 4). Hive control bees were at least 5 days younger than all other experimental groups, suggesting that age could also influence the behavioral discrepancies we saw in our study, as we know that experience plays a role in influencing fanning behavior in honeybees [52]. Regardless, our results suggest that a group’s ability to properly balance factors related to their social dynamics is essential for successfully producing a group behavior, lending support to our hypothesis.

Effective communication largely relies on the decision-making process and social behavior of all individuals within a group [1,20,66], which is heavily influenced by factors like group size and individual variation [19,66,71,72]. The rate, modularity, and feedback loops of information transfer grant animal groups the adaptability required to respond to changing environment conditions [4]. For example, variation in learning phenotypes in honeybees modulate colony foraging via differences in the enthusiastic performance of the waggle dance [73]. Leeches, locusts, and fruit flies will behave differently depending on the presence of others in their environment and prior in-group experience[74–76]. Thus, group members are constantly balancing whether and when to behave based on the information available and their own biological state, resulting in the maintenance of cohesiveness or group status [30–33]. A disruption in any of these components can lead to groups failing to work together to efficiently produce a collective behavior and respond to the environment.

Bees and other organisms frequently face human pollutants as an ongoing challenge in their environment [77–83]. Antibiotics, as persistent environmental pollutants spread by human activity [57,84–86], are particularly concerning, as they disrupt the fundamental physiology of many organisms [85,86]. Although life-saving for the treatment of infectious diseases, antibiotics can have ramifications on the individual outside of just gut dysbiosis [56,87–89]. For example, antibiotics can interact with the endocrine system [90], alter gene expression [56,91,92], paradoxically increase disease susceptibility [54,93], and disrupt social behavior by altering host physiology[43,87]. Such individualistic effects when considered on a much wider scale, like across whole animal groups, can potentially scale to perturb population dynamics[81]. In honeybees, the specific mechanisms through which antibiotics can cause behavioral and physiological detriment remain largely unclear. It is possible that antibiotics, outside of disrupting the gut microbiome, could destroy sensory structures important for social organization, as seen in schooling fish lateral hair cell studies [43,94]. These potential consequences have led recent FDA regulations on antibiotic use in the agricultural industry, which is the largest consumer of antibiotics in the United States [90,95,96]. Regardless, as humans continue to have lasting impacts on the global ecosystem [85,86], it is imperative to identify the mechanisms by which group-living organisms respond to and adapt to their environment within social contexts. Our results begin to explore this gap in knowledge by contributing key insight into how individual perturbation can cascade to ultimately affect group performance of a behavior.

Organisms, from single-celled bacteria to workers of a eusocial insect colony to members of a bird flock, engage in complex behavioral dynamics to respond to an ever-changing environment. The mechanisms that we’ve identified here surrounding social efficiency can apply to not only various animal systems but also human engineering, such as distributed robotics and computing [97,98]. Further pursuing the mechanisms behind collective behavior in animals can potentially unlock prospects for designing efficient algorithms and systems for human use. This work begins to unravel the social mechanisms that drive a collective dynamic and highlights the importance of efficiency in balancing tradeoffs associated with complex systems.

## Acknowledgments

We would like to thank Gabe Smith for his assistance in collecting data and Casey Lambert for her assistance in analyzing tracking data. We also would like to thank David Farynyk, the creator of ABCTracker, for his guidance on navigating and using his program for our study, and Daniel Charbonneau for providing the basic R code pipeline to analyze ABCTracker data. Thank you to members of the Cook lab for their helpful feedback on an early version of this manuscript, as well as the 2 reviewers and the editor that greatly improved the final version.

## Author Contribution

Conceptualization, J.B.N and C.N.C.; Methodology, J.B.N and C.N.C.; Validation, J.B.N.; Formal Analysis, J.B.N; Investigation, J.B.N.; Resources, C.N.C.; Data curation, J.B.N.; Writing – Original Draft, J.B.N and C.N.C.; Writing – Review & Editing, C.N.C.; Visualization, J.B.N; Supervision, C.N.C.; Project Administration, C.N.C.

## Supplementary Figures

**Table S1.**
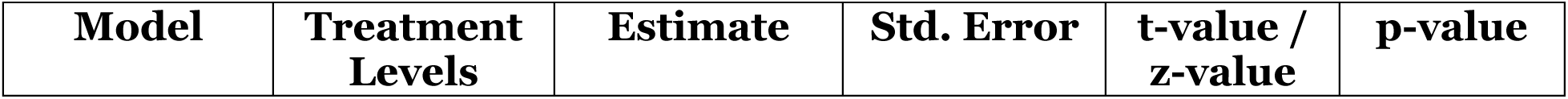
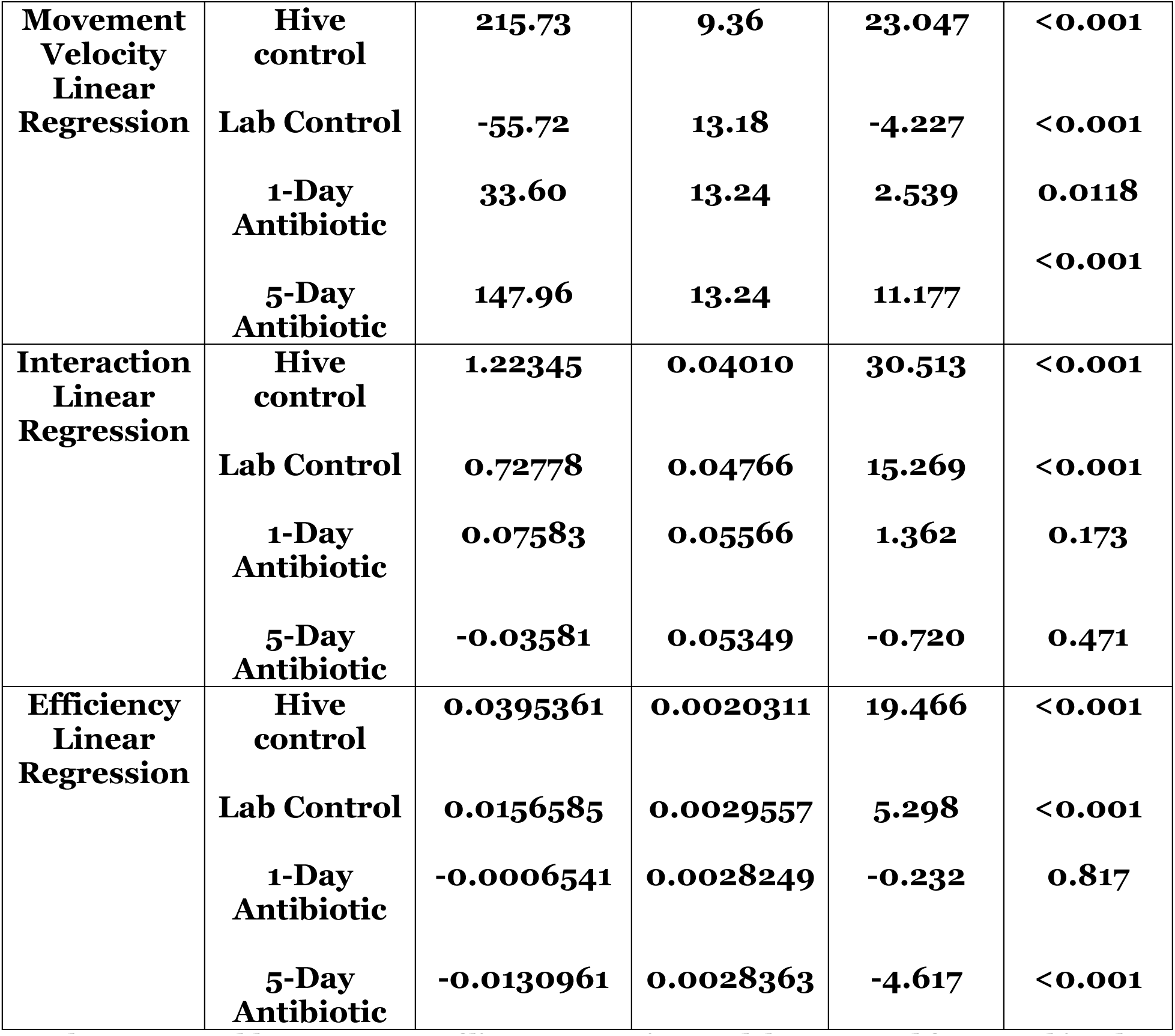
Summary of linear regression models generated from tracking data. For our movement velocity linear regression model, hive control, lab control, and the 5-day antibiotic treatments had a large significant effect on movement velocity. For our interaction linear regression model, the hive control and lab control group alone had a significant effect on the interactions bees had within videos. For our efficiency linear regression model, the hive control, lab control, and 5-day antibiotic group all had significant effects on the efficiency of interactions in our videos. These data demonstrate the effect sizes of our treatment groups in our study.

**Supplementary Figure 2.**
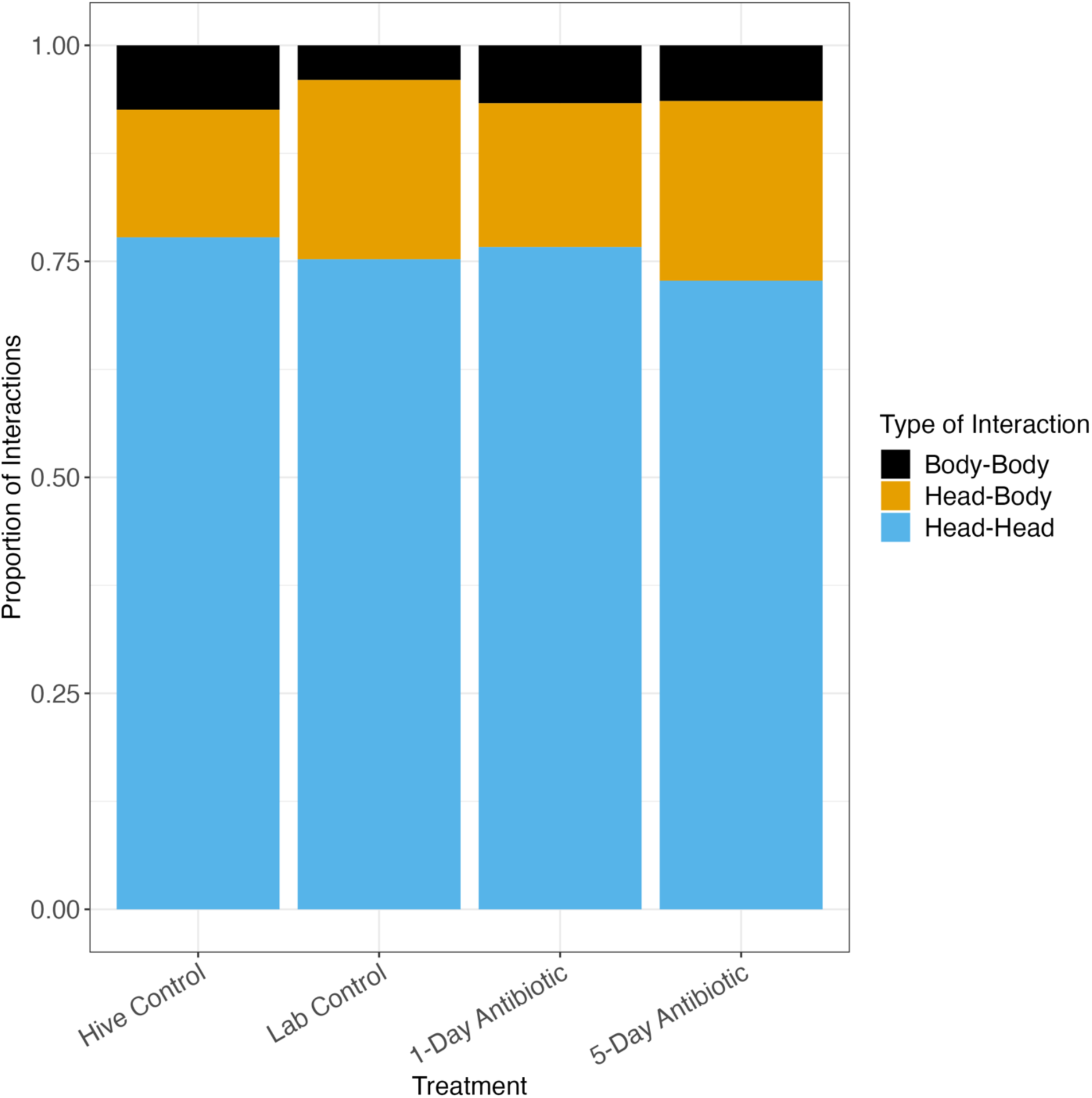
Head-to-head interactions dominate honeybee interaction type. Proportion of the types of interactions that occurred across all the interactions that happened within our videos. Head-to-head interactions accounted for 75% of all interactions across all treatment groups.

**Supplementary Figure 3.**
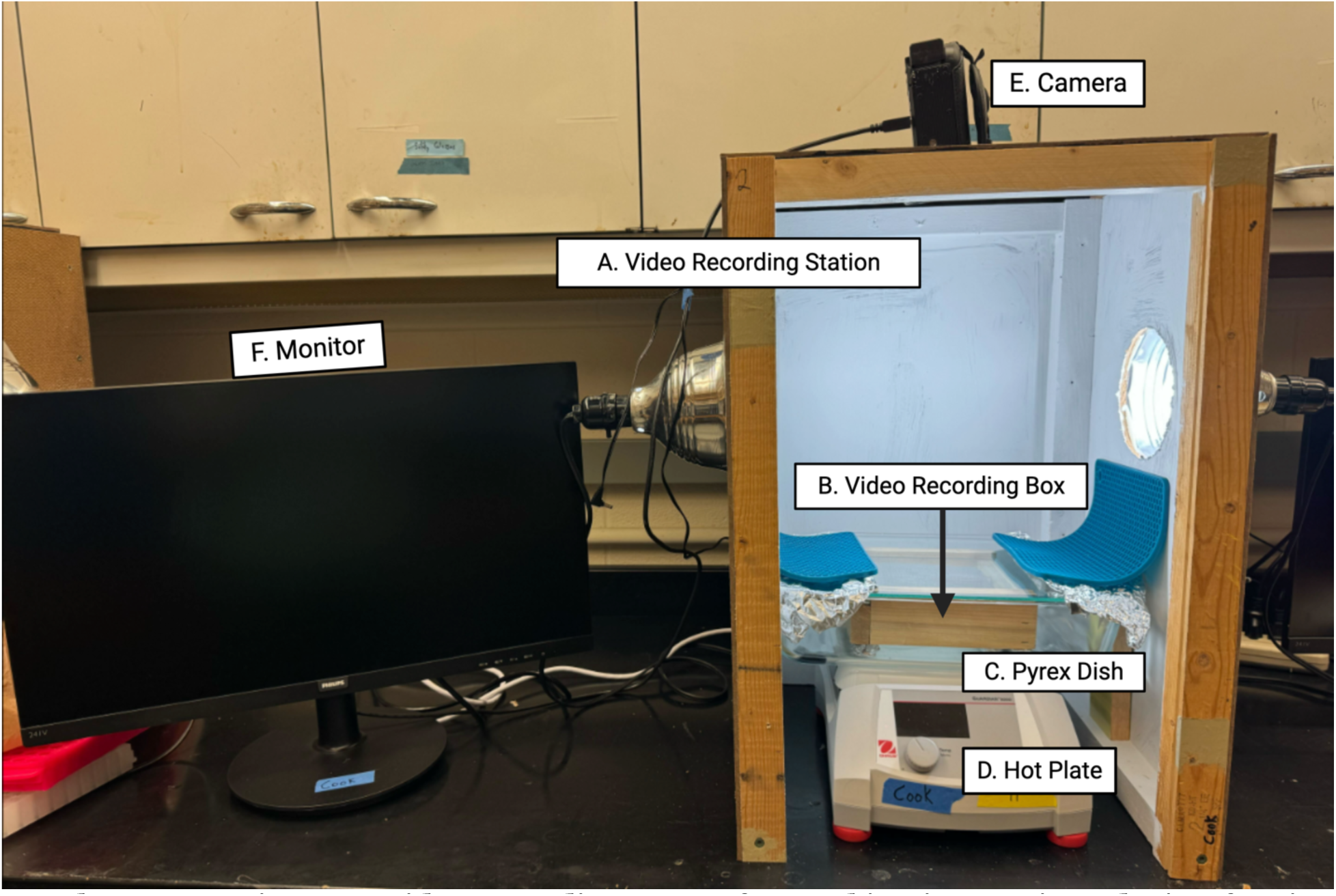
Video recording set-up for tracking interactions during fanning assay. Video recording station (A) was built using plywood and 2x4s with lamps affixed to cut holes on either side. Station was placed over a hot plate (D), and after marking, bees were housed in a wooden box of similar volume (B) to small fanning cages (354.98cm^3^). The bottom of the wooden box was lined with mesh wire and a silicone baking mat to minimize surface heat exposure to the bees, and the temperature probe was fed into the box spatially separated from where fanners were housed. This wooden box was then placed into a 13 x 8.5 x 3.25-inch glass baking dish (Pyrex) with a glass lid on top (C) to mimic the fanning assay set up. Aluminum foil was used to cover any holes, and silicone mats were used to cover the foil to minimize camera glare. A Panasonic HC-V800 camera (E) was then placed at the top of the station and was connected to a computer monitor to allow for experimenters to observe the bees (F). Once video recording set up was completed, a plywood cover was slotted over the open front of the station just above the hot plate to minimize reflections in the glass. This video recording station gave a top-down, clear view of the bees during the duration of a fanny assay while minimizing 3D space and subsequent video analysis difficulties. Figure was created using Biorender.

**Supplementary Figure 4.**
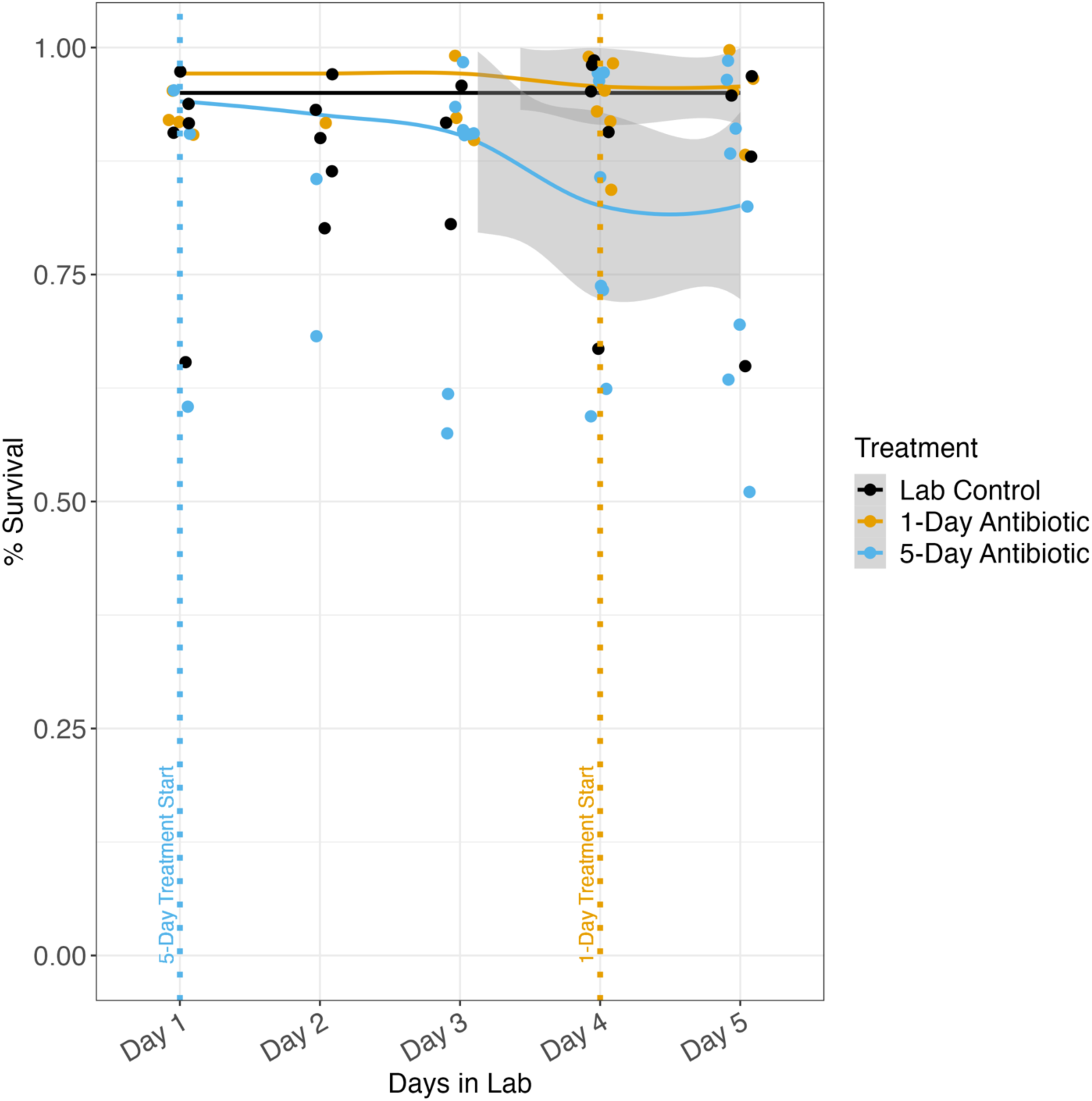
2022 preliminary data for honeybee survival during treatment duration. Survival curves for the duration of our experimental treatment period. All bees were housed in groups of 30 consisting of the same colony members. nlab control= 8, n1-day= 7, n5-day= 7, for n = boxes of bees. Survival curves calculated for the duration of treatment period in 2022. No difference was found in survival. Logistic regression, X^2^ = 0.64147, p = 0.7256. Shaded regions indicate 95% confidence intervals.

**Supplementary Figure 5.**
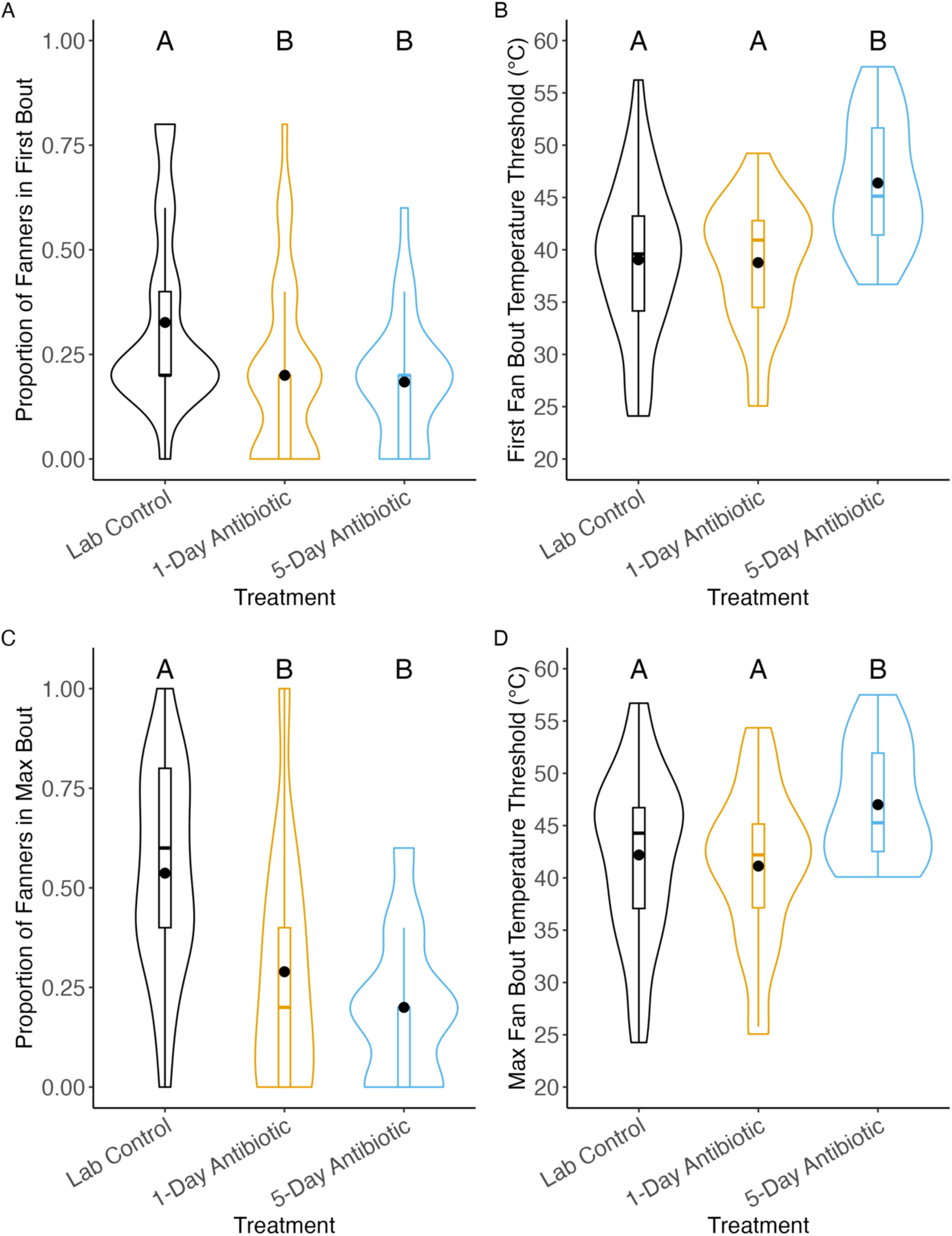
2022 preliminary data for oxytetracycline negatively impacting fanning behavior in honeybees. Fanner honeybees subjected to oxytetracycline treatment experience a deficit in their fanning behavior in comparison to bees not treated with antibiotics. Lab control = bees exposed to a 40% sucrose solution for 5 days, 1-day = bees exposed to oxytetracycline 24 hours prior to fanning assay, and 5-day = bees exposed to oxytetracycline for 5 days. n_lab control_= 38, n_1-day_ = 38, n_5-day_ = 25, for n = cage of 5 bees. A) Honeybees that were treated with oxytetracycline, regardless of time exposed, are significantly less likely to be part of the first fanning bout than bees that were not treated B) Honeybees that fanned in the first bout did so at significantly higher temperatures C) Honeybees that were treated with oxytetracycline, regardless of time exposed, are significantly less likely to fan during the max bout of fanning than those that were not. Logistic regression, X^2^=44.08, p<0.001. D) Fanners treated with oxytetracycline for 5 days fanned at significantly higher temperatures during the max bout of fanning than those not treated with oxytetracycline and those treated for 24 hours prior. Linear regression, F= 8.289, p = 0.0006.

**Supplementary Figure 6.**
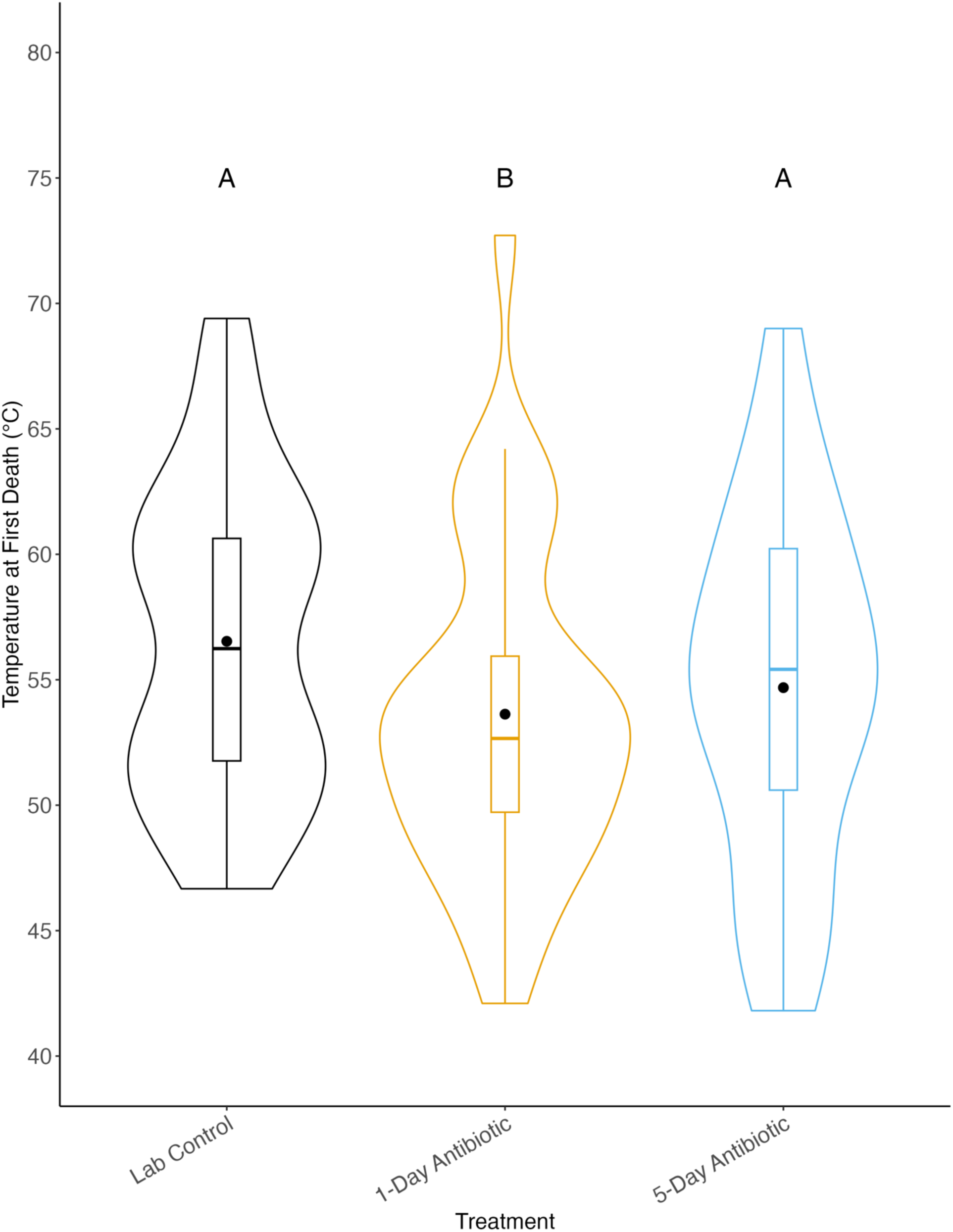
2022 preliminary data for antibiotic treatment lowering temperature at which first death occurs during fanning assay. Middle dots indicate mean, middle lines indicate median. Letters indicate statistical significance. Linear regression, F=6.0976, p<0.001, Tukey post-hoc test, p<0.05. nlab control = 38, n1-day = 38, n5-day = 25, for n = cage of 5 bees.

## References

1. Couzin ID. Collective cognition in animal groups. Trends in Cognitive Sciences. 2009;13: 36–43. doi:10.1016/j.tics.2008.10.002

2. Gordon DM. The Ecology of Collective Behavior. PLOS Biology. 2014;12: e1001805. doi:10.1371/journal.pbio.1001805

3. Ouellette NT, Gordon DM. Goals and Limitations of Modeling Collective Behavior in Biological Systems. Frontiers in Physics. 2021;9. Available: https://www.frontiersin.org/articles/10.3389/fphy.2021.687823

4. Gordon DM. Collective behavior in relation with changing environments: Dynamics, modularity, and agency. Evolution & Development. 2023;25: 430–438. doi:10.1111/ede.12439

5. Jolles JW, Boogert NJ, Sridhar VH, Couzin ID, Manica A. Consistent Individual Differences Drive Collective Behavior and Group Functioning of Schooling Fish. Curr Biol. 2017;27: 2862–2868.e7. doi:10.1016/j.cub.2017.08.004

6. Schaerf TM, Dillingham PW, Ward AJW. The effects of external cues on individual and collective behavior of shoaling fish. Science Advances. 2017;3: e1603201. doi:10.1126/sciadv.1603201

7. Ward, Sumpter DJT, Couzin ID, Hart PJB, Krause J. Quorum decision-making facilitates information transfer in fish shoals. Proceedings of the National Academy of Sciences. 2008;105: 6948–6953. doi:10.1073/pnas.0710344105

8. Ward, Krause J, Sumpter DJT. Quorum Decision-Making in Foraging Fish Shoals. PLOS ONE. 2012;7: e32411. doi:10.1371/journal.pone.0032411

9. Davidson JD, Gordon DM. Spatial organization and interactions of harvester ants during foraging activity. Journal of The Royal Society Interface. 2017;14: 20170413. doi:10.1098/rsif.2017.0413

10. Pagliara R, Gordon DM, Leonard NE. Regulation of harvester ant foraging as a closed-loop excitable system. PLOS Computational Biology. 2018;14: e1006200. doi:10.1371/journal.pcbi.1006200

11. Yusufaly TI, Boedicker JQ. Mapping quorum sensing onto neural networks to understand collective decision making in heterogeneous microbial communities. Phys Biol. 2017;14: 046002. doi:10.1088/1478-3975/aa7c1e

12. Vasconcelos MM, Mitra U, Camara O, Silva KP, Boedicker J. Bacterial Quorum Sensing as a Networked Decision System. 2018 IEEE International Conference on Communications (ICC). Kansas City, MO: IEEE; 2018. pp. 1–6. doi:10.1109/ICC.2018.8422668

13. Bridges AA, Bassler BL. The intragenus and interspecies quorum-sensing autoinducers exert distinct control over Vibrio cholerae biofilm formation and dispersal. PLOS Biology. 2019;17: e3000429. doi:10.1371/journal.pbio.3000429

14. Krause J, Krause P of FB and EJ, Ruxton GD, Ruxton G, Ruxton IG. Living in Groups. OUP Oxford; 2002.

15. Majolo B, de Bortoli Vizioli A, Schino G. Costs and benefits of group living in primates: group size effects on behaviour and demography. Animal Behaviour. 2008;76: 1235–1247. doi:10.1016/j.anbehav.2008.06.008

16. Ward A, Webster M. Sociality: The Behaviour of Group-Living Animals. Cham: Springer International Publishing; 2016. doi:10.1007/978-3-319-28585-6

17. Bergman TJ, Beehner JC. Information Ecology: an integrative framework for studying animal behavior. Trends in Ecology & Evolution. 2023;38: 1041–1050. doi:10.1016/j.tree.2023.05.017

18. Ioannou CC, Laskowski KL. A multi-scale review of the dynamics of collective behaviour: from rapid responses to ontogeny and evolution. Philosophical Transactions of the Royal Society B: Biological Sciences. 2023;378: 20220059. doi:10.1098/rstb.2022.0059

19. Pacala SW, Gordon DM, Godfray HCJ. Effects of social group size on information transfer and task allocation. Evol Ecol. 1996;10: 127–165. doi:10.1007/BF01241782

20. Couzin ID, Krause J. Self-Organization and Collective Behavior in Vertebrates. Advances in the Study of Behavior. Elsevier; 2003. pp. 1–75. doi:10.1016/S0065-3454(03)01001-5

21. Falcón-Cortés A, Boyer D, Ramos-Fernández G. Collective learning from individual experiences and information transfer during group foraging. Journal of The Royal Society Interface. 2019;16: 20180803. doi:10.1098/rsif.2018.0803

22. Pratt SC. Behavioral mechanisms of collective nest-site choice by the ant Temnothorax curvispinosus. Insect Soc. 2005;52: 383–392. doi:10.1007/s00040-005-0823-z

23. Berdahl A, Torney CJ, Ioannou CC, Faria JJ, Couzin ID. Emergent Sensing of Complex Environments by Mobile Animal Groups. Science. 2013;339: 574–576. doi:10.1126/science.1225883

24. Seeley TD, Visscher PK. Quorum sensing during nest-site selection by honeybee swarms. Behav Ecol Sociobiol. 2004;56: 594–601. doi:10.1007/s00265-004-0814-5

25. Visscher PK. Group Decision Making in Nest-Site Selection Among Social Insects. Annual Review of Entomology. 2007;52: 255–275. doi:10.1146/annurev.ento.51.110104.151025

26. Mglich M, Maschwitz U, Hölldobler B. Tandem Calling: A New Kind of Signal in Ant Communication. Science. 1974;186: 1046–1047. doi:10.1126/science.186.4168.1046

27. Franklin EL, Franks. Individual and social learning in tandem-running recruitment by ants. Animal Behaviour. 2012;84: 361–368. doi:10.1016/j.anbehav.2012.05.002

28. Grüter C, Farina WM. The honeybee waggle dance: can we follow the steps? Trends in Ecology & Evolution. 2009;24: 242–247. doi:10.1016/j.tree.2008.12.007

29. Al Toufailia H, Couvillon MJ, Ratnieks FLW, Grüter C. Honey bee waggle dance communication: signal meaning and signal noise affect dance follower behaviour. Behav Ecol Sociobiol. 2013;67: 549–556. doi:10.1007/s00265-012-1474-5

30. Franks, Dornhaus A, Fitzsimmons JP, Stevens M. Speed versus accuracy in collective decision making. Proceedings of the Royal Society of London Series B: Biological Sciences. 2003;270: 2457–2463. doi:10.1098/rspb.2003.2527

31. Chittka L, Skorupski P, Raine NE. Speed–accuracy tradeoffs in animal decision making. Trends in Ecology & Evolution. 2009;24: 400–407. doi:10.1016/j.tree.2009.02.010

32. Franks, Richardson TO, Stroeymeyt N, Kirby RW, Amos WMD, Hogan PM, et al. Speed– cohesion trade-offs in collective decision making in ants and the concept of precision in animal behaviour. Animal Behaviour. 2013;85: 1233–1244. doi:10.1016/j.anbehav.2013.03.010

33. Heitz RP. The speed-accuracy tradeoff: history, physiology, methodology, and behavior. Frontiers in Neuroscience. 2014;8. Available: https://www.frontiersin.org/articles/10.3389/fnins.2014.00150

34. Greene MJ, Gordon DM. Interaction rate informs harvester ant task decisions. Behavioral Ecology. 2007;18: 451–455. doi:10.1093/beheco/arl105

35. Pinter-Wollman N, Bala A, Merrell A, Queirolo J, Stumpe MC, Holmes S, et al. Harvester ants use interactions to regulate forager activation and availability. Animal Behaviour. 2013;86: 197–207. doi:10.1016/j.anbehav.2013.05.012

36. Granovetter M. Threshold Models of Collective Behavior. American Journal of Sociology. 1978;83: 1420–1443. doi:10.1086/226707

37. Theraulaz G, Bonabeau E, Denuebourg J-N. Response threshold reinforcements and division of labour in insect societies. Proceedings of the Royal Society of London Series B: Biological Sciences. 1998;265: 327–332. doi:10.1098/rspb.1998.0299

38. Sakata H, Katayama N. Ant defence system: A mechanism organizing individual responses into efficient collective behavior. Ecol Res. 2001;16: 395–403. doi:10.1046/j.1440-1703.2001.00404.x

39. Koolhaas JM, Korte SM, De Boer SF, Van Der Vegt BJ, Van Reenen CG, Hopster H, et al. Coping styles in animals: current status in behavior and stress-physiology. Neuroscience & Biobehavioral Reviews. 1999;23: 925–935. doi:10.1016/S0149-7634(99)00026-3

40. Wright GA, Lillvis JL, Bray HJ, Mustard JA. Physiological State Influences the Social Interactions of Two Honeybee Nest Mates. PLOS ONE. 2012;7: e32677. doi:10.1371/journal.pone.0032677

41. Gordon DM. The Ecology of Collective Behavior in Ants. 2018.

42. Faucher K, Parmentier E, Becco C, Vandewalle N, Vandewalle P. Fish lateral system is required for accurate control of shoaling behaviour. Animal Behaviour. 2010;79: 679–687. doi:10.1016/j.anbehav.2009.12.020

43. Suli A, Watson GM, Rubel EW, Raible DW. Rheotaxis in Larval Zebrafish Is Mediated by Lateral Line Mechanosensory Hair Cells. PLOS ONE. 2012;7: e29727. doi:10.1371/journal.pone.0029727

44. Anderson M. The Evolution of Eusociality. Annual Review of Ecology and Systematics. 1984;15: 165–189. doi:10.1146/annurev.es.15.110184.001121

45. Anderson C, Ratnieks FLW. Worker allocation in insect societies: coordination of nectar foragers and nectar receivers in honey bee (Apis mellifera) colonies. Behav Ecol Sociobiol. 1999;46: 73–81. doi:10.1007/s002650050595

46. Tarpy DR, Gilley DC, Seeley TD. Levels of selection in a social insect: a review of conflict and cooperation during honey bee (Apis mellifera) queen replacement. Behav Ecol Sociobiol. 2004;55: 513–523. doi:10.1007/s00265-003-0738-5

47. Calderone NW. Insect Pollinated Crops, Insect Pollinators and US Agriculture: Trend Analysis of Aggregate Data for the Period 1992–2009. PLOS ONE. 2012;7: e37235. doi:10.1371/journal.pone.0037235

48. Hung K-LJ, Kingston JM, Albrecht M, Holway DA, Kohn JR. The worldwide importance of honey bees as pollinators in natural habitats. Proceedings of the Royal Society B: Biological Sciences. 2018;285: 20172140. doi:10.1098/rspb.2017.2140

49. Lindauer M. The Water Economy and Temperature Regulation of the Honeybee Colony. Bee World. 1955;36: 81–92. doi:10.1080/0005772X.1955.11094876

50. Egley RL, Breed MD. The Fanner Honey Bee: Behavioral Variability and Environmental Cues in Workers Performing a Specialized Task. J Insect Behav. 2013;26: 238–245. doi:10.1007/s10905-012-9357-1

51. Cook CN, Breed MD. Social context influences the initiation and threshold of thermoregulatory behaviour in honeybees. Animal Behaviour. 2013;86: 323–329. doi:10.1016/j.anbehav.2013.05.021

52. Kaspar RE, Cook CN, Breed MD. Experienced individuals influence the thermoregulatory fanning behaviour in honey bee colonies. Animal Behaviour. 2018;142: 69–76. doi:10.1016/j.anbehav.2018.06.004

53. Evans JD. Diverse origins of tetracycline resistance in the honey bee bacterial pathogen *Paenibacillus larvae*. Journal of Invertebrate Pathology. 2003;83: 46–50. doi:10.1016/S0022-2011(03)00039-9

54. Raymann K, Shaffer Z, Moran NA. Antibiotic exposure perturbs the gut microbiota and elevates mortality in honeybees. PLoS Biol. 2017;15. doi:10.1371/journal.pbio.2001861

55. Ortiz-Alvarado Y, Clark DR, Vega-Melendez CJ, Flores-Cruz Z, Domingez-Bello MG, Giray T. Antibiotics in hives and their effects on honey bee physiology and behavioral development. Biology Open. 2020;9. doi:10.1242/bio.053884

56. Ortiz-Alvarado Y, Giray T. Antibiotics Alter the Expression of Genes Related to Behavioral Development in Honey Bees (Hymenoptera: Apidae). Journal of Insect Science. 2022;22: 10. doi:10.1093/jisesa/ieac017

57. Wang X, Zhang L, Han C, Zhang Y, Zhuo J. Simulation study of oxytetracycline contamination remediation in groundwater circulation wells enhanced by nano-calcium peroxide and ozone. Sci Rep. 2023;13: 9136. doi:10.1038/s41598-023-36310-1

58. Seeley TD. Adaptive significance of the age polyethism schedule in honeybee colonies. Behav Ecol Sociobiol. 1982;11: 287–293. doi:10.1007/BF00299306

59. Winston ML, Punnett EN. Factors determining temporal division of labor in honeybees. Can J Zool. 1982;60: 2947–2952. doi:10.1139/z82-372

60. Richards ED, Tell LA, Davis JL, Baynes RE, Lin Z, Maunsell FP, et al. Honey bee medicine for veterinarians and guidance for avoiding violative chemical residues in honey. Journal of the American Veterinary Medical Association. 2021;259: 860–873. doi:10.2460/javma.259.8.860

61. Masood F, Thebeau JM, Cloet A, Kozii IV, Zabrodski MW, Biganski S, et al. Evaluating approved and alternative treatments against an oxytetracycline-resistant bacterium responsible for European foulbrood disease in honey bees. Sci Rep. 2022;12: 5906. doi:10.1038/s41598-022-09796-4

62. Pettis JS, Kochansky J, Feldlaufer MF. Larval Apis mellifera L. (Hymenoptera: Apidae) Mortality After Topical Application of Antibiotics and Dusts. Journal of Economic Entomology. 2004;97: 171–176. doi:10.1093/jee/97.2.171

63. 63. Rice L. A beginning-to-end system for efficiently gathering tracking data on multiple targets. Doctoral dissertation, The University of North Carolina at Charlotte.; 2016.

64. Rice L, Tate S, Farynyk D, Sun J, Chism G, Charbonneau D, et al. ABCTracker: an easy-to-use, cloud-based application for tracking multiple objects. arXiv; 2020. doi:10.48550/arXiv.2001.10072

65. Wild B, Dormagen DM, Zachariae A, Smith ML, Traynor KS, Brockmann D, et al. Social networks predict the life and death of honey bees. Nat Commun. 2021;12: 1110. doi:10.1038/s41467-021-21212-5

66. Kashetsky T, Yan J, Doering G, Skelton T, Dukas R. The effect of experience on collective decision-making. Behavioural Processes. 2023;213: 104962. doi:10.1016/j.beproc.2023.104962

67. Martel A-C, Zeggane S, Drajnudel P, Faucon J-P, Aubert M. Tetracycline residues in honey after hive treatment. Food Additives & Contaminants. 2006;23: 265–273. doi:10.1080/02652030500469048

68. Currie S, LeBlanc S, Watters MA, Gilmour KM. Agonistic encounters and cellular angst: social interactions induce heat shock proteins in juvenile salmonid fish. Proceedings of the Royal Society B: Biological Sciences. 2009;277: 905–913. doi:10.1098/rspb.2009.1562

69. Uchida S, Hara K, Kobayashi A, Fujimoto M, Otsuki K, Yamagata H, et al. Impaired hippocampal spinogenesis and neurogenesis and altered affective behavior in mice lacking heat shock factor 1. Proceedings of the National Academy of Sciences. 2011;108: 1681– 1686. doi:10.1073/pnas.1016424108

70. Yusishen ME, Yoon GR, Bugg W, Jeffries KM, Currie S, Andersono WG. Love thy neighbor: Social buffering following exposure to an acute thermal stressor in a gregarious fish, the lake sturgeon (*Acipenser fulvescens*). Comparative Biochemistry and Physiology Part A: Molecular & Integrative Physiology. 2020;243: 110686. doi:10.1016/j.cbpa.2020.110686

71. King AJ, Cowlishaw G. When to use social information: the advantage of large group size in individual decision making. Biology Letters. 2007;3: 137–139. doi:10.1098/rsbl.2007.0017

72. Sokolowski MB. Social Interactions in “Simple” Model Systems. Neuron. 2010;65: 780–794. doi:10.1016/j.neuron.2010.03.007

73. Cook CN, Lemanski NJ, Mosqueiro T, Ozturk C, Gadau J, Pinter-Wollman N, et al. Individual learning phenotypes drive collective behavior. Proc Natl Acad Sci USA. 2020;117: 17949. doi:10.1073/pnas.1920554117

74. Bisson G, Torre V. Statistical characterization of social interactions and collective behavior in medicinal leeches. Journal of Neurophysiology. 2011;106: 78–90. doi:10.1152/jn.01043.2010

75. Knebel D, Sha-ked C, Agmon N, Ariel G, Ayali A. Collective motion as a distinct behavioral state of the individual. iScience. 2021;24: 102299. doi:10.1016/j.isci.2021.102299

76. Bentzur A, Ben-Shaanan S, Benichou JIC, Costi E, Levi M, Ilany A, et al. Early Life Experience Shapes Male Behavior and Social Networks in Drosophila. Current Biology. 2021;31: 486–501.e3. doi:10.1016/j.cub.2020.10.060

77. Travis CC, Hester ST. Global chemical pollution. Environ Sci Technol. 1991;25: 814–819. doi:10.1021/es00017a001

78. Dachs J, Méjanelle L. Organic Pollutants in Coastal Waters, Sediments, and Biota: A Relevant Driver for Ecosystems During the Anthropocene? Estuaries and Coasts. 2010;33: 1–14. doi:10.1007/s12237-009-9255-8

79. Marcantonio R, Javeline D, Field S, Fuentes A. Global distribution and coincidence of pollution, climate impacts, and health risk in the Anthropocene. PLOS ONE. 2021;16: e0254060. doi:10.1371/journal.pone.0254060

80. Hill MK. Understanding Environmental Pollution. Cambridge University Press; 2020.

81. Schell LM, Denham M. Environmental Pollution in Urban Environments and Human Biology. Annual Review of Anthropology. 2003;32: 111–134. doi:10.1146/annurev.anthro.32.061002.093218

82. McArt SH, Fersch AA, Milano NJ, Truitt LL, Böröczky K. High pesticide risk to honey bees despite low focal crop pollen collection during pollination of a mass blooming crop. Sci Rep. 2017;7: 46554. doi:10.1038/srep46554

83. Martinello M, Manzinello C, Dainese N, Giuliato I, Gallina A, Mutinelli F. The Honey Bee: An Active Biosampler of Environmental Pollution and a Possible Warning Biomarker for Human Health. Applied Sciences. 2021;11: 6481. doi:10.3390/app11146481

84. Aminov R. A Brief History of the Antibiotic Era: Lessons Learned and Challenges for the Future. Frontiers in Microbiology. 2010;1. Available: https://www.frontiersin.org/articles/10.3389/fmicb.2010.00134

85. Gothwal R, Shashidhar T. Antibiotic Pollution in the Environment: A Review. CLEAN – Soil, Air, Water. 2015;43: 479–489. doi:10.1002/clen.201300989

86. Kraemer SA, Ramachandran A, Perron GG. Antibiotic Pollution in the Environment: From Microbial Ecology to Public Policy. Microorganisms. 2019;7: 180. doi:10.3390/microorganisms7060180

87. Wu W-L, Adame MD, Liou C-W, Barlow JT, Lai T-T, Sharon G, et al. Microbiota regulate social behaviour via stress response neurons in the brain. Nature. 2021;595: 409–414. doi:10.1038/s41586-021-03669-y

88. Hayer SS, Hwang S, Clayton JB. Antibiotic-induced gut dysbiosis and cognitive, emotional, and behavioral changes in rodents: a systematic review and meta-analysis. Front Neurosci. 2023;17: 1237177. doi:10.3389/fnins.2023.1237177

89. Ramirez J, Guarner F, Bustos Fernandez L, Maruy A, Sdepanian VL, Cohen H. Antibiotics as Major Disruptors of Gut Microbiota. Front Cell Infect Microbiol. 2020;10. doi:10.3389/fcimb.2020.572912

90. Dibner JJ, Richards JD. Antibiotic growth promoters in agriculture: history and mode of action. Poultry Science. 2005;84: 634–643. doi:10.1093/ps/84.4.634

91. Ward TL, Weber BP, Mendoza KM, Danzeisen JL, Llop K, Lang K, et al. Antibiotics and Host-Tailored Probiotics Similarly Modulate Effects on the Developing Avian Microbiome, Mycobiome, and Host Gene Expression. mBio. 2019;10: 10.1128/mbio.02171-19. doi:10.1128/mbio.02171-19

92. Sun L, Zhang X, Zhang Y, Zheng K, Xiang Q, Chen N, et al. Antibiotic-Induced Disruption of Gut Microbiota Alters Local Metabolomes and Immune Responses. Front Cell Infect Microbiol. 2019;9. doi:10.3389/fcimb.2019.00099

93. Keeney KM, Yurist-Doutsch S, Arrieta M-C, Finlay BB. Effects of Antibiotics on Human Microbiota and Subsequent Disease. Annual Review of Microbiology. 2014;68: 217–235. doi:10.1146/annurev-micro-091313-103456

94. Harris JA, Cheng AG, Cunningham LL, MacDonald G, Raible DW, Rubel EW. Neomycin-induced hair cell death and rapid regeneration in the lateral line of zebrafish (Danio rerio). J Assoc Res Otolaryngol. 2003;4: 219–234. doi:10.1007/s10162-002-3022-x

95. Kumar K, C. Gupta S, Chander Y, Singh AK. Antibiotic Use in Agriculture and Its Impact on the Terrestrial Environment. Advances in Agronomy. Academic Press; 2005. pp. 1–54. doi:10.1016/S0065-2113(05)87001-4

96. Mann A, Nehra K, Rana JS, Dahiya T. Antibiotic resistance in agriculture: Perspectives on upcoming strategies to overcome upsurge in resistance. Current Research in Microbial Sciences. 2021;2: 100030. doi:10.1016/j.crmicr.2021.100030

97. Brambilla M, Ferrante E, Birattari M, Dorigo M. Swarm robotics: a review from the swarm engineering perspective. Swarm Intell. 2013;7: 1–41. doi:10.1007/s11721-012-0075-2

98. Kar AK. Bio inspired computing – A review of algorithms and scope of applications. Expert Systems with Applications. 2016;59: 20–32. doi:10.1016/j.eswa.2016.04.018

