## Supplemental Figures for "Disruption of Collective Behavior Correlates with Reduced Interaction Efficiency"

### Supplementary Figures

| Model | Treatment Levels | Estimate | Std. Error | t-value / z-value | p-value |
|---|---|---|---|---|---|
| Movement Velocity Linear Regression | Hive control | 215.73 | 9.36 | 23.047 | <0.001 |
| | Lab Control | -55.72 | 13.18 | -4.227 | <0.001 |
| | 1-Day Antibiotic | 33.60 | 13.24 | 2.539 | 0.0118 |
| | 5-Day Antibiotic | 147.96 | 13.24 | 11.177 | <0.001 |
| Interaction Linear Regression | Hive control | 1.22345 | 0.04010 | 30.513 | <0.001 |
| | Lab Control | 0.72778 | 0.04766 | 15.269 | <0.001 |
| | 1-Day Antibiotic | 0.07583 | 0.05566 | 1.362 | 0.173 |
| | 5-Day Antibiotic | -0.03581 | 0.05349 | -0.720 | 0.471 |
| Efficiency Linear Regression | Hive control | 0.0395361 | 0.0020311 | 19.466 | <0.001 |
| | Lab Control | 0.0156585 | 0.0029557 | 5.298 | <0.001 |
| | 1-Day Antibiotic | -0.0006541 | 0.0028249 | -0.232 | 0.817 |
| | 5-Day Antibiotic | -0.0130961 | 0.0028363 | -4.617 | <0.001 |

**Supplementary Table 1. Summary of linear regression models generated from tracking data.** For our movement velocity linear regression model, hive control, lab control, and the 5-day antibiotic treatments had a large significant effect on movement velocity. For our interaction linear regression model, the hive control and lab control group alone had a significant effect on the interactions bees had within videos. For our efficiency linear regression model, the hive control, lab control, and 5-day antibiotic group all had significant effects on the efficiency of interactions in our videos. These data demonstrate the effect sizes of our treatment groups in our study.

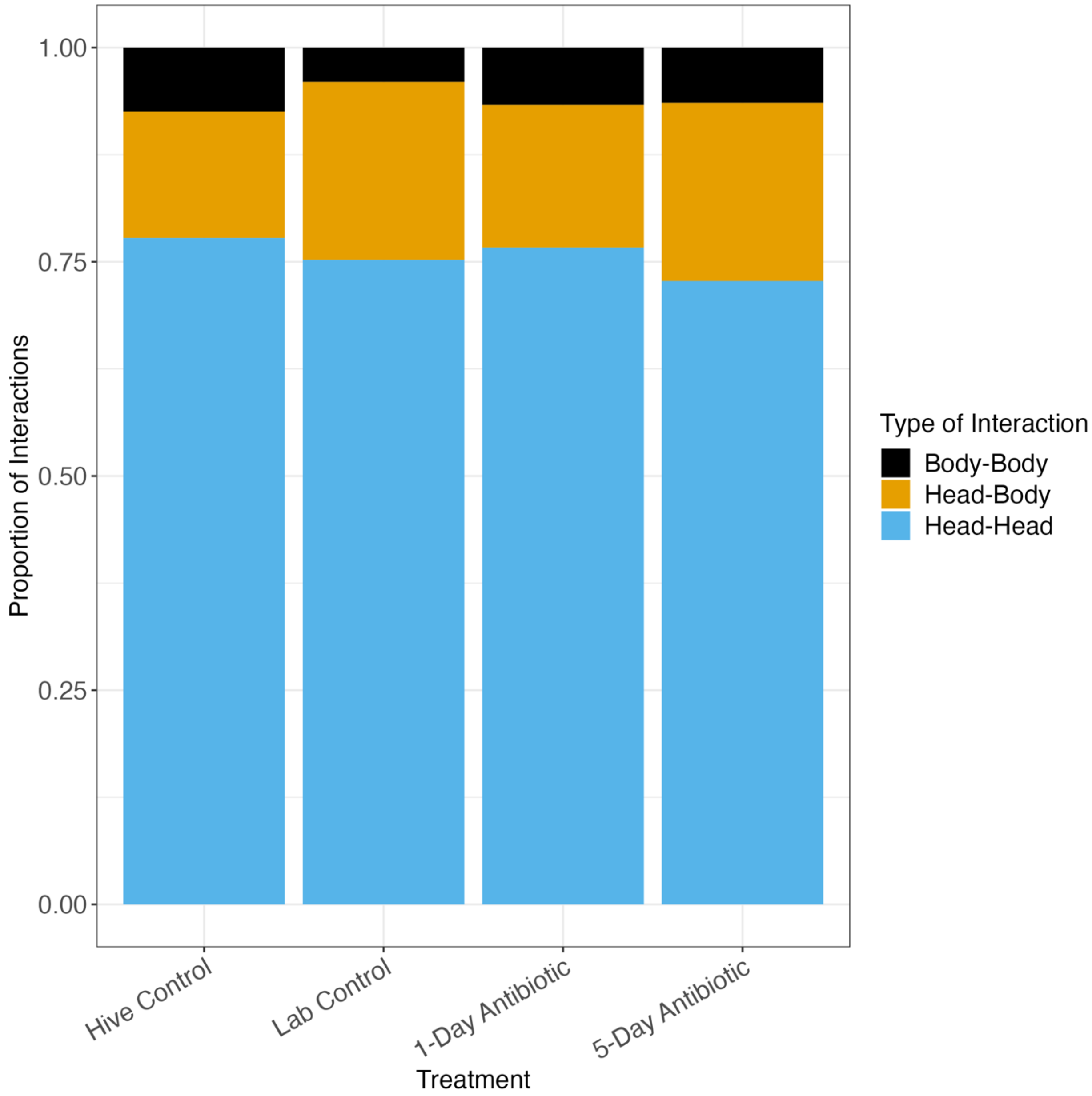


**Supplementary Figure 2. Head-to-head interactions dominate honeybee interaction type.** Proportion of the types of interactions that occurred across all the interactions that happened within our videos. Head-to-head interactions accounted for 75% of all interactions across all treatment groups.

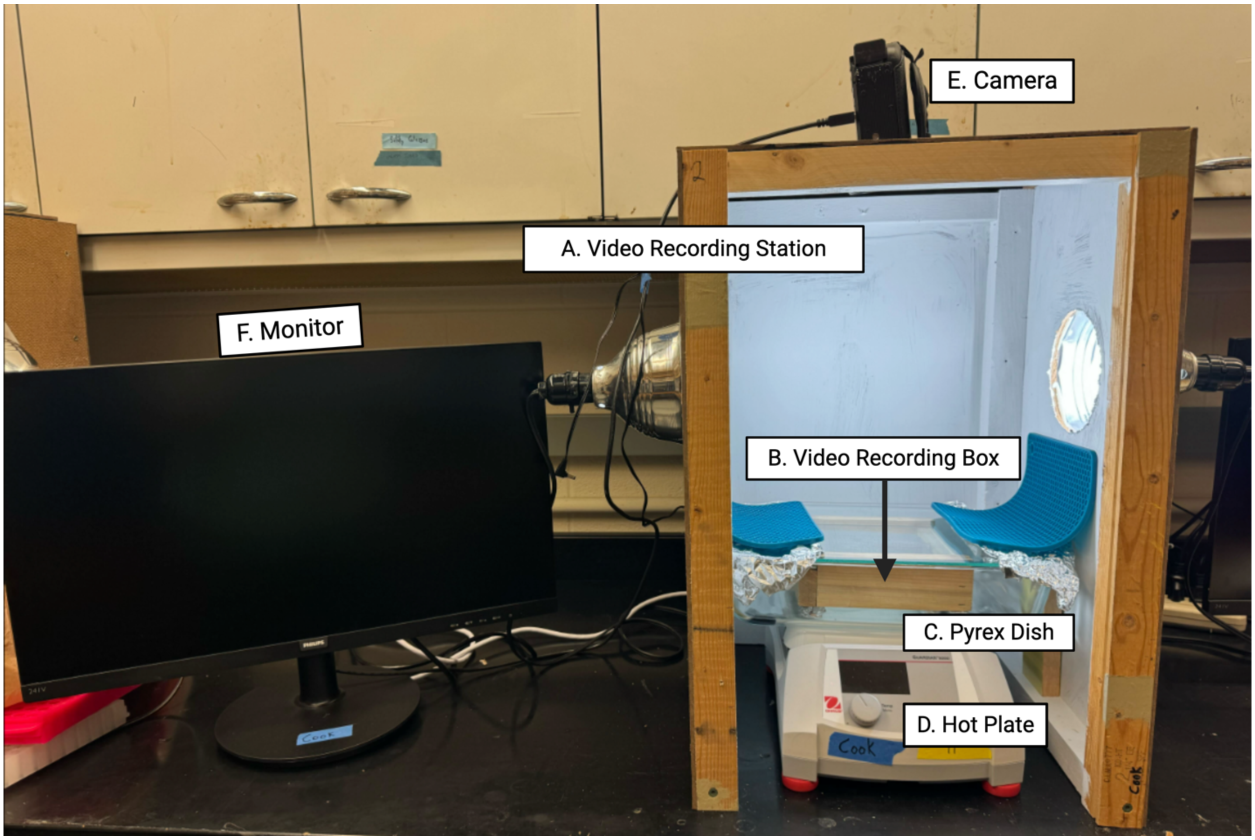


**Supplementary Figure 3. Video recording set-up for tracking interactions during fanning assay.** Video recording station (A) was built using plywood and 2x4s with lamps affixed to cut holes on either side. Station was placed over a hot plate (D), and after marking, bees were housed in a wooden box of similar volume (B) to small fanning cages (354.98$cm^3$). The bottom of the wooden box was lined with mesh wire and a silicone baking mat to minimize surface heat exposure to the bees, and the temperature probe was fed into the box spatially separated from where fanners were housed. This wooden box was then placed into a 13 x 8.5 x 3.25-inch glass baking dish (Pyrex) with a glass lid on top (C) to mimic the fanning assay set up. Aluminum foil was used to cover any holes, and silicone mats were used to cover the foil to minimize camera glare. A Panasonic HC-V800 camera (E) was then placed at the top of the station and was connected to a computer monitor to allow for experimenters to observe the bees (F). Once video recording set up was completed, a plywood cover was slotted over the open front of the station just above the hot plate to minimize reflections in the glass. This video recording station gave a top-down, clear view of the bees during the duration of a fanny assay while minimizing 3D space and subsequent video analysis difficulties. Figure was created using Biorender.

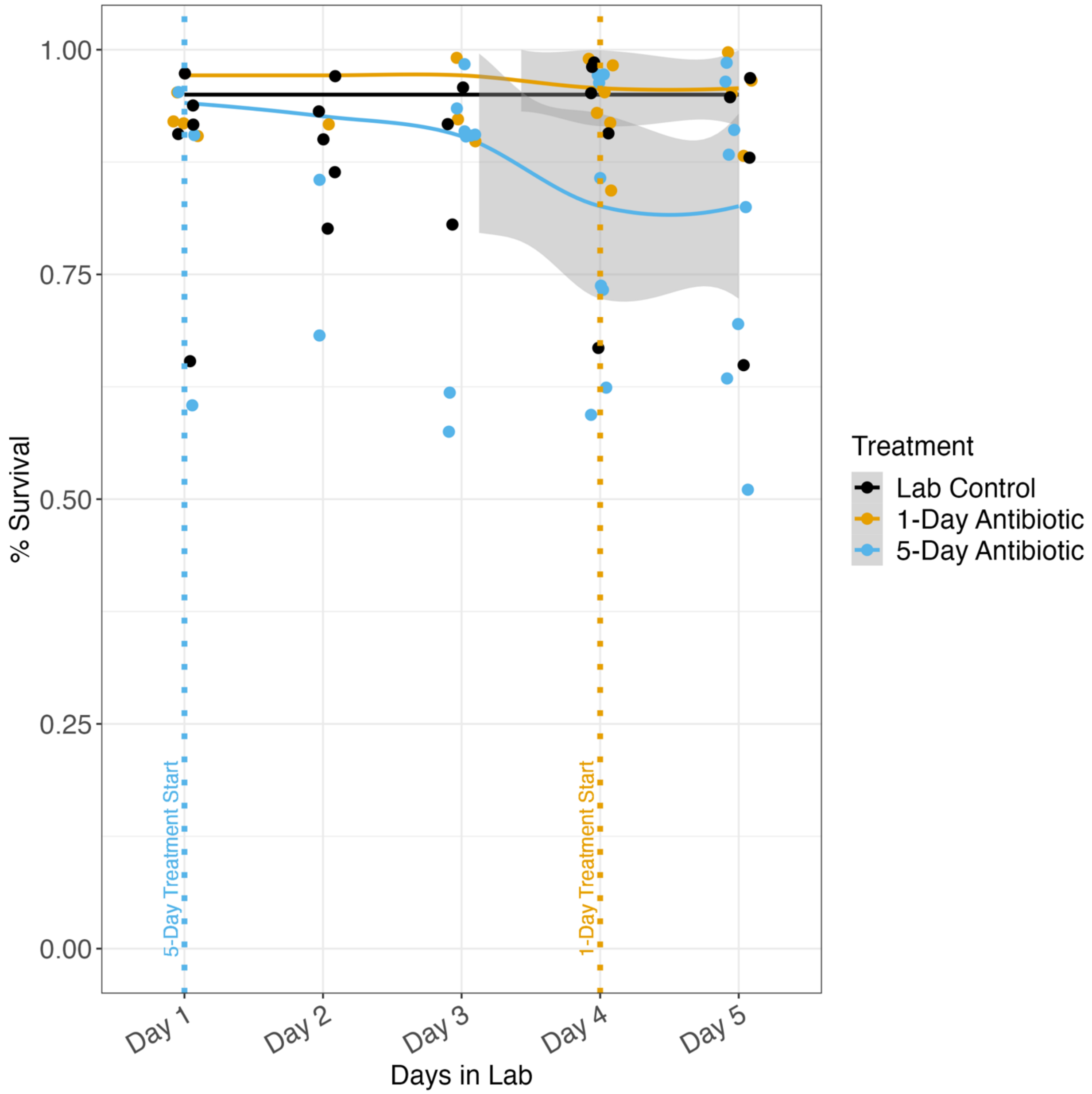


**Supplementary Figure 4. 2022 preliminary data for honeybee survival during treatment duration.** Survival curves for the duration of our experimental treatment period. All bees were housed in groups of 30 consisting of the same colony members. $n_{lab\ control}$= 8, $n_{1\text{-}day}$= 7, $n_{5\text{-}day}$= 7, for n = boxes of bees. Survival curves calculated for the duration of treatment period in 2022. No difference was found in survival. Logistic regression, $X^2$ = 0.64147, p = 0.7256. Shaded regions indicate 95% confidence intervals.

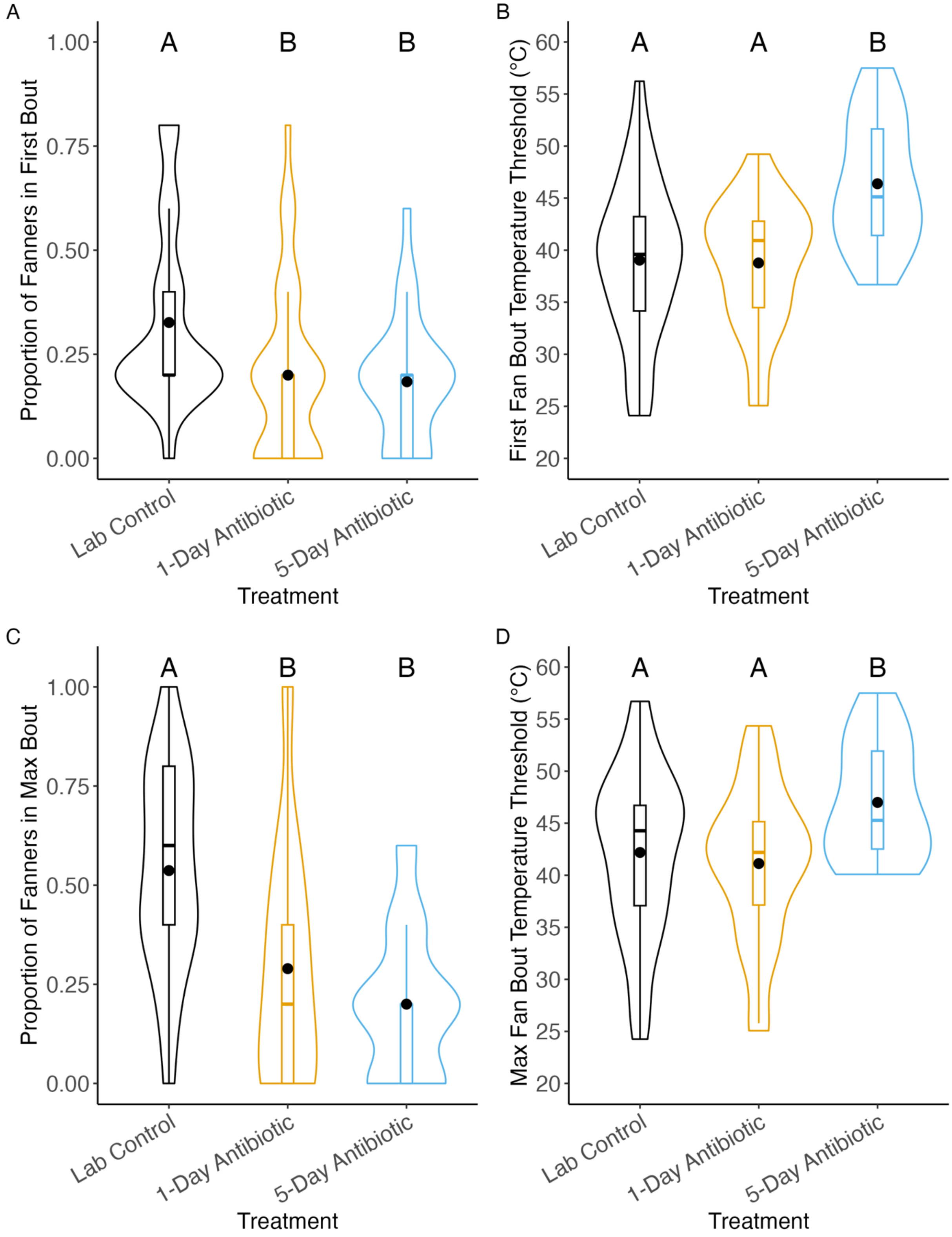


**Supplementary Figure 5. 2022 preliminary data for oxytetracycline negatively impacting fanning behavior in honeybees.** Fanner honeybees subjected to oxytetracycline treatment experience a deficit in their fanning behavior in comparison to bees not treated with antibiotics. Lab control = bees exposed to a 40% sucrose solution for 5 days, 1-day = bees exposed to oxytetracycline 24 hours prior to fanning assay, and 5-day = bees exposed to oxytetracycline for 5 days. $n_{lab\ control}$= 38, $n_{1\text{-}day}$ = 38, $n_{5\text{-}day}$ = 25, for n = cage of 5 bees. A) Honeybees that were treated with oxytetracycline, regardless of time exposed, are significantly less likely to be part of the first fanning bout than bees that were not treated B) Honeybees that fanned in the first bout did so at significantly higher temperatures C) Honeybees that were treated with oxytetracycline, regardless of time exposed, are significantly less likely to fan during the max bout of fanning than those that were not. Logistic regression, $X^2$=44.08, $p<0.001$. D) Fanners treated with oxytetracycline for 5 days fanned at significantly higher temperatures during the max bout of fanning than those not treated with oxytetracycline and those treated for 24 hours prior. Linear regression, F= 8.289, p = 0.0006.

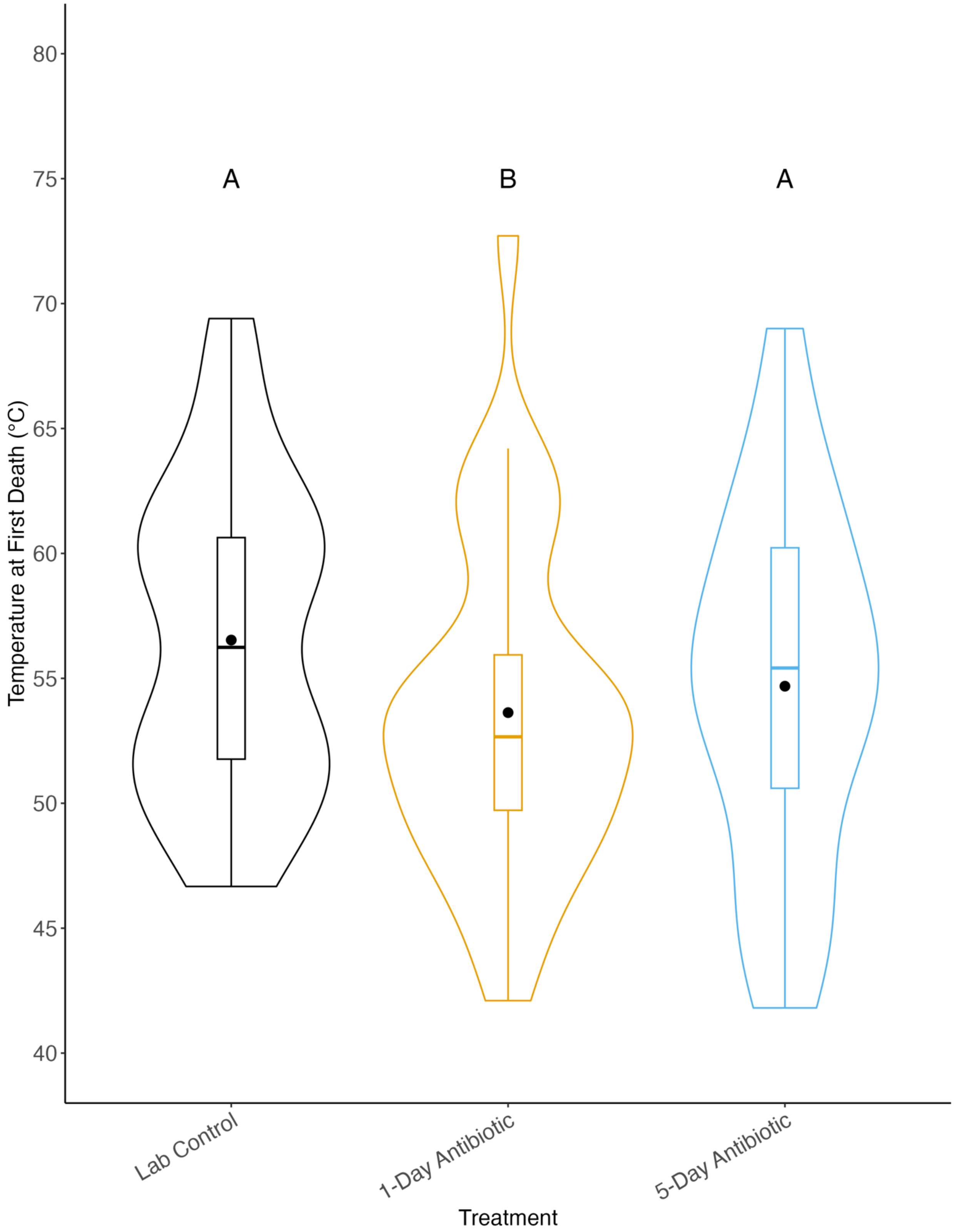


**Supplementary Figure 6. 2022 preliminary data for antibiotic treatment lowering temperature at which first death occurs during fanning assay.** Middle dots indicate mean, middle lines indicate median. Letters indicate statistical significance. Linear regression, $F=6.0976$, $p<0.001$, Tukey post-hoc test, $p<0.05$. $n_{lab\ control} = 38$, $n_{1\text{-}day} = 38$, $n_{5\text{-}day} = 25$, for n = cage of 5 bees.
